# Analysis of the Cross-Study Replicability of Tuberculosis Gene Signatures Using 49 Curated Transcriptomic Datasets

**DOI:** 10.1101/2023.12.01.569442

**Authors:** Xutao Wang, Katie Harper, Pranay Sinha, W. Evan Johnson, Prasad Patil

## Abstract

**Background:** Tuberculosis (TB) is the leading cause of infectious disease mortality worldwide. Numerous blood-based gene expression signatures have been proposed in the literature as alternative tools for diagnosing TB infection. Ongoing efforts are actively focused on developing additional signatures in other TB-related contexts. However, the generalizability of these signatures to different patient contexts is not well-characterized. There is a pressing need for a well-curated database of TB gene expression studies for the systematic assessment of existing and newly developed TB gene signatures.

**Results:** We built the curatedTBData, a manually-curated database of 49 TB transcriptomic studies. This data resource is freely available through GitHub and as an R Bioconductor package that allows users to validate new and existing biomarkers without the challenges of harmonizing heterogeneous studies. We also demonstrate the use of this data resource with cross-study comparisons for 72 TB gene signatures. For the comparison of subjects with active TB from healthy controls, 19 gene signatures had weighted mean AUC of 0.90 or greater, with the highest result of 0.94. In active TB disease versus latent TB infection, 7 gene signatures had weighted mean AUC of 0.90 or greater, with a maximum of 0.93. We also explore ensembling methods for averaging predictions from multiple gene signatures to significantly improve diagnostic ability beyond any single signature.

**Conclusions:** The curatedTBData data package offers a comprehensive resource of curated gene expression and clinically annotated data. It could be used to identify robust new TB gene signatures, to perform comparative analysis of existing TB gene signatures, and to develop alternative gene set scoring or ensembling methods, among other things. This resource will also facilitate the development of new signatures that are generalizable across cohorts or more applicable to specific subsets of patients (e.g. with rare comorbid conditions, etc.). We demonstrated that these blood-based gene signatures could distinguish patients with distinct TB outcomes; moreover, the combination of multiple gene signatures could improve the overall predictive accuracy in differentiating these subtypes, which point out an important aspect for the translation of genomics to clinical implementation.

## Introduction

Approximately 1.8 billion people in the world are estimated to be carriers the the causative pathogen for TB, *Mycobacterium tuberculosis*, often denoted as latent TB infection (LTBI), and about 10 million people develop ‘active’ pulmonary TB (PTB) each year worldwide (1). Sputum culture has been widely used for TB diagnosis, but it usually takes 6–7 days for a positive diagnosis and up to 42 days for a confirmed negative diagnosis (2); furthermore, sputum polymerase chain reaction (PCR) has become more common and offers faster results, but is expensive and unable to detect extrapulmonary TB. More recently, the use of GenXpert MTB/RIF has increased the speed and accuracy of diagnostic; the remaining challenges in TB diagnosis focus around paucibacillary TB (acid fast bacilli smear negative pulmonary TB, extra-pulmonary TB, and pediatric TB) and among individuals with certain comorbid conditions. As such, researchers have attempted to address these last diagnostic needs using blood-based gene expression signatures. For example, these have been developed for distinguishing active TB from LTBI, other bacterial and viral infections, and from healthy controls with high accuracy (among other applications). However, studies used to develop these signatures often represent different geographical regions, distinct age groups, and cohorts with varying comorbid conditions (2; 3; 4). As a result of this study heterogeneity, a single TB signature may exhibit poor generalizability in diverse patient populations (5; 6). Variation in statistical procedures and over-fitting of training models used to develop these signatures may also contribute to reduced generalizability (2; 7; 8).

As more TB transcriptomic datasets become publicly available, there is greater opportunity to identify more robust and generalizable gene signatures for TB outcomes in diverse cohorts (9). However, differences in gene expression platforms, annotation, pre-processing, and batch effects across studies often prevents users from making efficient use of these resources (10). As a result, there is no current database of TB gene expression studies that provides researchers harmonized datasets for gene signature development and validation.

Here, we introduce the curatedTBData, an R data package available through GitHub and Bioconductor. This package enhances the reproducibility and replicability of signature development and validation, facilitating integrative-analyses across TB gene expression datasets. Currently, the curatedTBData provides harmonized gene expression measurements and clinical annotations for 49 studies. These data are stored in R asMultiassayExperiment objects (11), which efficiently manages the raw data, multiple normalized versions of the data, metadata, and some intermediate analysis results. We have developed harmonized data preprocessing and annotation pipelines, and reprocessed each of the 49 datasets (Figure 1). The curatedTBData also provides functions for subsetting and aggregating the transcriptomic datasets using existing tools, such as ComBat and ComBat-Seq among other tools (12; 13; 14). To demonstrate the utility of the curatedTBData, we conducted a systematic comparison and validation of multiple published blood-based TB gene signatures (15). We examine the predictive ability of 72 TB gene signatures across the 49 curated studies in the context of differentiating PTB from healthy controls (Control) and PTB from LTBI. We also use dataset and datatype aggregation tools based on our previous work (16) and built into curatedTBData to train and evaluate ensembles of gene signatures.

**Figure 1.**
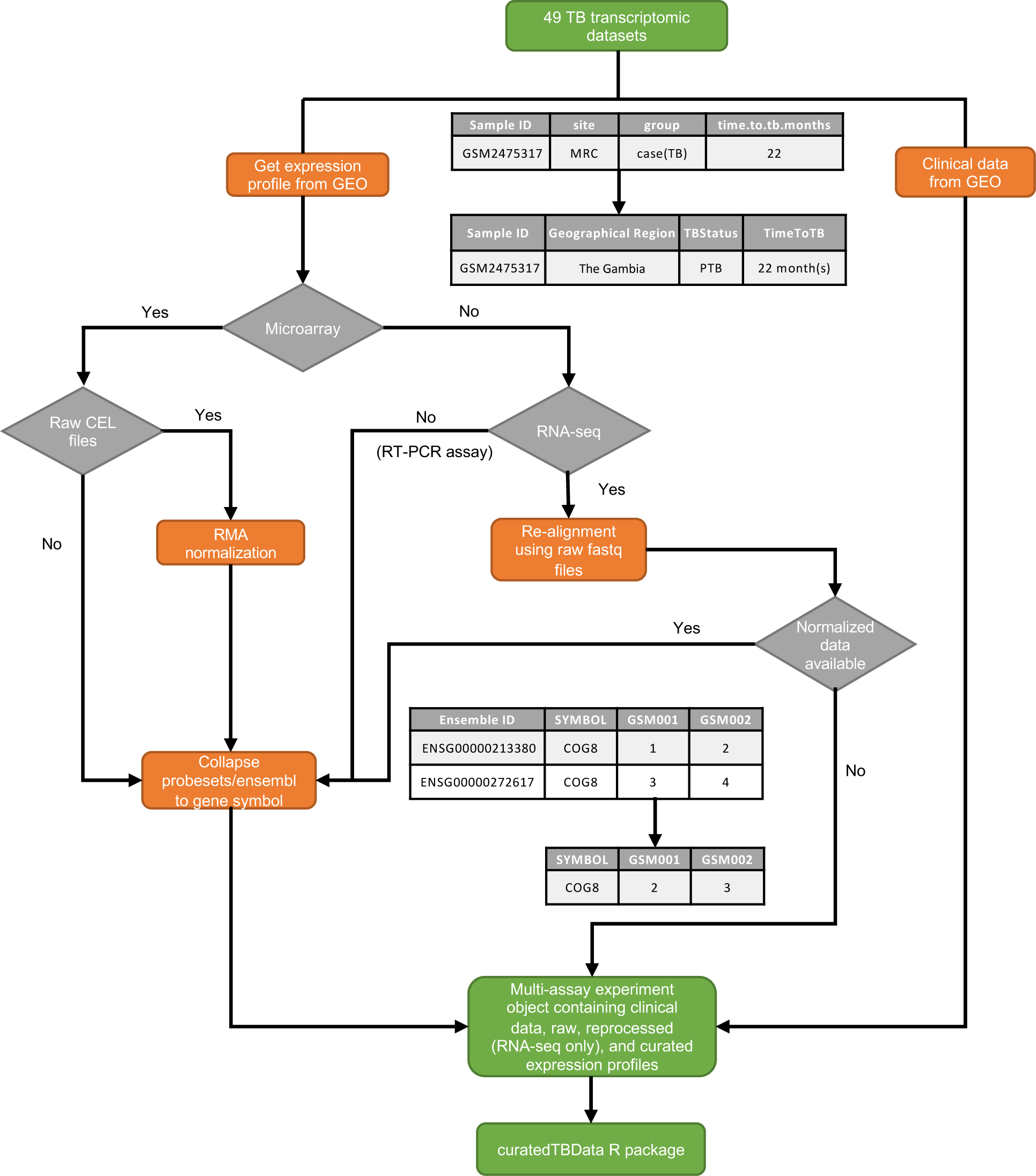
Flowchart of processing steps for curating TB transcriptomic datasets.

In this paper, we describe details of the curatedTBData data curation process. We provide examples in the comparison of TB gene signatures using gene set scoring methods. Furthermore, we showcase the application of an ensemble learning strategy, which combines the predictive ability from multiple gene signatures.

## Methods and Implementation

### Study selection

We selected our studies using a step-wise process. Initially, priority was given to datasets that were used as training sets for signatures that are present in our TBSignatureProfiler R package (15). Of the 49 datasets, 34 of them match this criteria. In addition, there are 11 datasets (679 subjects) that correspond to TBSignatureProfiler biomarker training sets that we are actively curating. Finally, 49 studies were curated for convenience for our prior studies (8; 6). Curation of these datasets is an active, ongoing process, especially for additional biomarker training datasets. After identifying the datasets, we employed the R package GEOquery (17) to download them from the Gene Expression Omnibus (GEO). The 49 TB transcriptome datasets in the textttcuratedTBData resource span 17 gene measurement platforms and contain clinical information for 4161 samples (Table S1). In order to access these datasets in a uniform way and to enhance the replicability of results across studies, we reprocessed these 49 datasets with manually curated clinical metadata information as described in Figure 1. For each transcriptomic study, both curated and raw data were included in the final output, and its reprocessing pipeline (in the form of R script) and related functions were also provided in the package (and in the Methods below).

### Metadata curation

For each study, we computationally and/or manually extracted the experimental, clinical, and demographic metadata by leveraging functions from the GEOquery (17) package. The curated metadata information was stored in the form of DataFrame, with standardized column names representing the different features (TBStatus, Gender, Age etc.) across different studies, and row names corresponding to each sample’s official GEO name. Moreover, the information within each feature was also aligned across datasets, so that users could efficiently subset samples based on certain criteria for meta-analysis. Finally, a standardized key of 60 variables to clinical and demographic information was summarized for current version of the curatedTBData (Table S2).

### Uniform preprocesing of RNA-sequencing data

We reprocessed each RNA sequencing dataset from the raw FASTQ files. The downloaded sequence data were first trimmed using Trimamomatic-0.36 (18) to remove low quality reads (SLIDINGWINDOW=4:20, LEADING=3, TRAILING=3, MINLEN=36). The trimmed data were then aligned to NCBI RefSeq gene annotations hg38 (for all RNA-seq studies) and hg19 (for selected RNA-seq studies where hg19 was used in their original paper); lastly, the mapped reads for genomics features were counted using the package Rsubread (19; 20). The pipeline of reprocessing RNA-Seq datasets can be found at the GitHub page for paper, and. We validated the reprocessed results by plotting the correlation between the reprocessed RNA-Seq data and its corresponding original data from GEO.

### Data storage and functions in the package

In the package, the S4 object MultiAssayExperiment (11) was implemented to store all the information for each study. This data structure is beneficial for representing and analyzing multi-omics experiments. The curatedTBData package takes the advantage of using the MultiAssayExperiment to retain raw expression profiles, normalized expression profiles, and different versions of the data etc. In such case, researchers can efficiently manage and access both raw and multiple processed/harmonized version of each study. Additionally, object of MultiAssayExperiment from the curatedTBData can be converted to SummarizedExperiment object easily, enabling users to leverage toolkit from other packages (i.e. TBSignatureProfiler).

The curatedTBData provides analytic utility functions for biomarker validation along with comparative visualizations including, boxplot, heatmap, and rideplot. Additionally, it includes function (subset_curatedTBData) that subsets samples based on input condition and function (combine_object) that facilitates the integration of datasets.

### Datasets and gene signatures used for comparison

Currently, there are 49 transcriptomic studies included in curatedTBData. For demonstration purpose, we compared the performance of 72 for their content and metadata for differentiating patients with PTB from Control or from LTBI. Study selection/merging and sample selection process was conducted before the comparative analysis (Figure 2). Specifically, datasets with only PTB subjects or that focus on progression analysis were excluded (n = 8), as well as subjects with HIV, diabetes, and involved in repeated measurement other than baseline were excluded. Finally, 19 studies that compared patients with PTB to patients with LTBI and 15 studies that compared patients with PTB to healthy controls had been included for downstream comparative analysis.

**Figure 2.**
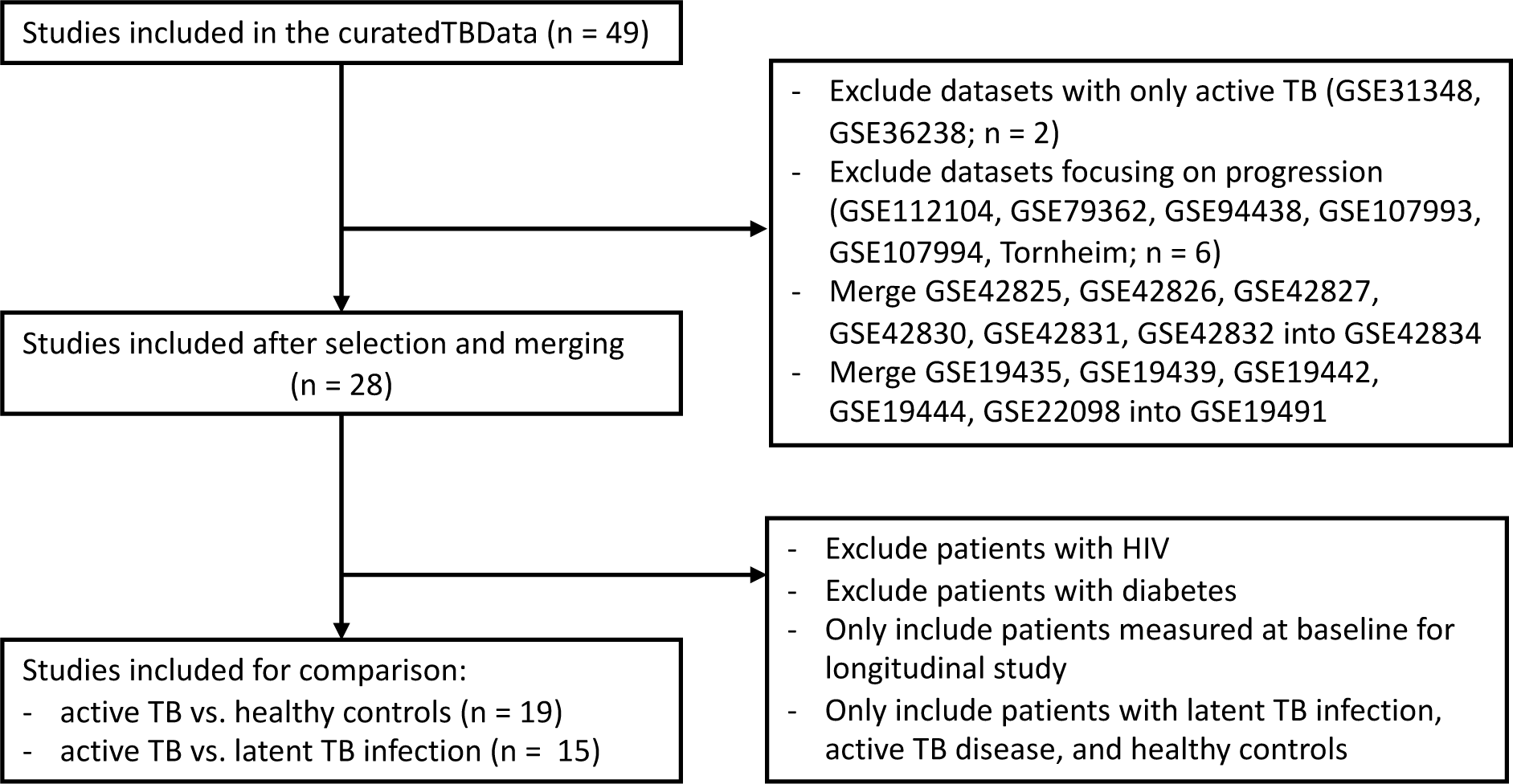
Flowchart of datasets selection process for comparative analysis.

Because the gene sets which define each model are often more readily available than the original signature model itself, we chose to perform the comparative analysis using two uniform single-sample methods (21): 1. Single-Sample Gene Set Enrichment Analysis (ssGSEA) (22) and 2. Pathway Level Analysis of Gene Expression (PLAGE) (23), which represent two competitive and self-contained gene set scoring methods (24). ssGSEA calculates a sample-level gene set score by comparing the distribution of gene expression ranks inside and outside the gene set (22); whereas PLAGE standardizes expression values and applies singular value decomposition (SVD) for the genes within the gene set. The activity level of a pathway in a given sample is taken as the coefficient of the first right-singular vector (25). The R package GSVA (26) was implemented to compute scores from ssGSEA and PLAGE. There are chances that some of genes within gene sets were missing across multiple transcriptomic datasets, primarily due to different versions for the name of the gene symbols or the inconsistent gene coverage across sequencing platforms. To alleviate this problem, the R package HGNCHelper (27) was used to update gene symbols for both TB gene signatures and transcriptomic datasets.

### Cross-study ensemble learning

Integrating information from different biomarkers is one of the critical applications of the curatedTBData database (16). In the context of ensemble stacking strategy (28), We viewed multiple gene signatures as one predictor known as ‘meta-predictor’ and trained ensemble learning models base on multiple studies to evaluate the the performance of this composite predictor.

As a demonstration exercise, 34 gene signatures has been selected in the example case of usage, and the list of signatures for training an ensemble model are shown in Figure S4. Additionally, datasets with sample size less than 15 were excluded from ensemble analysis. These datasets frequently exhibited strong signal among different TB subtypes, with a majority of biomarkers demonstrating high predictive accuracy for them. Finally, a total of 14 datasets for PTB vs. Control and 14 datasets for PTB vs. LTBI had been included in the example.

The procedures for performing ensemble analysis were listed as follows. Firstly, a subset of signatures with size *S* were randomly sampled without replacement to investigate how ensemble process varies across different gene sets (*S* = 5 in this example). Consider *K* training datasets and *V* validation datasets of dimension *n_k_* × *S*, where *k* = 1, 2*,…, K* + *V* denotes the dataset index and *S* represents the number of gene signatures applied to each dataset (i.e. each dataset has *S* columns of gene signature risk scores). Within each dataset we have an outcome (TB subtype or control) ***Y_k_*** and sets of predictors [***X*_1_*_,k_***, ***X*_2_*_,k_****,…,* ***X_S,k_***], where ***X_s,k_*** represents the predicted score given by gene signature *s* computed from either ssGSEA or PLAGE. Our final goal was to make predictions in the the validation datasets *V* using a linear combination of the predicted results for each of gene set. We define the predicted score obtained from training data *k* to be ***Ŷ*** *_k_* (***X_s,k_***) and the stacked predicted score from group of *K* datasets for gene set *s* to be ***Ŷ*** (***X_s_***), and the predicted results to a set of combined gene sets as 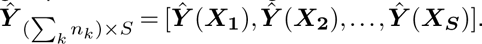. We determine weights *w_s_* that correspond to a set of combined gene signature scores based on the Lasso where

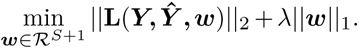

The final predicted score for the meta-predictor on validation dataset *v* is given by

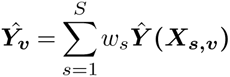

The performance of the ensemble sets on each validation study was evaluated using AUC, and the its overall validation performance was determined by the weighted mean AUC values (AUC*_w_*) across *V* validation studies:

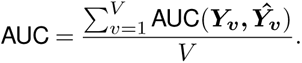

The variation of the AUCs for both ensemble sets and gene signature across *n* studies, each with observations: [*w*_1_*, w*_2_*,…, w_n_*] and AUCs: [*AU C*_1_*, AU C*_2_*,…, AU C_n_*], were evaluated using the weighted standard deviation (*SD_w_*) and showed in the form of ridge plot (Figure S6):

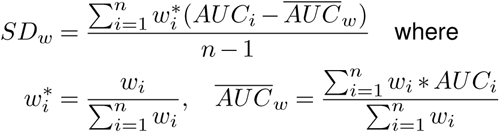

To reduce the bias of test studies selection, we performed cross-study validation by repeatedly splitting the multi-studies into *K* training and *V* testing datasets for *M* iterations. For the group of datasets with size *N* (*N* = 14 for both comparisons in this example), we randomly selected 80% of studies as training data with *K* = ⌊0.8*N* ⌋ and the remaining as testing data with *V* = *N* − *K* for *M* = 300 iterations.

### Statistics used for evaluation

For each TB gene signature, its performance in distinguishing different phenotype was be evaluated by the AUC (Area Under the Receiver Operating Characteristics Curve), sensitivity, and specificity value, calculated by the R package pROC (29). The sample-size weighted average AUC values (weighted AUCs), sensitivity values (weighted sensitivity), and specificity values (weighted specificity) were computed across datasets to evaluate the overall performance for each gene signature. The 95% confidence interval for these metrics was calculated over 1000 bootstrapped subsampling. Finally, the performance of ensemble of group signature was also compared with its single best subset using the weighted student’s t-test, implemented by the function wtd.t.test from R package weights (30).

### Availability of data and materials

The datasets generated are available in the: https://bioconductor.org/packages/release/data/ experiment/html/curatedTBData.html

The code for generating the results can be found in the: https://github.com/xutao-wang/curatedTBData_ paper_code

## Results

### Overall Performance of TB Gene Signatures

For the comparison of PTB vs. Control, three gene signatures (Blankley_5, Kaul_3, and Tabone_RES_27) out of 72 achieved the highest weighted AUCs of 0.94 (95% CI: 0.87 - 0.98) when assessed using ssGSEA; Conversely, another set of three gene signatures (Maertzdorf_15, Suliman_4, and Tabone_OD_11) had the highest weighed AUCs of 0.94 (95% CI: 0.88 - 0.98) using PLAGE (Table S3). For the comparison of PTB vs. LTBI, Bloom_RES_268, Estevez_133, and Tabone_RES_27 had the highest weighted AUCs of 0.92 (95% CI: 0.84 - 0.98) using ssGSEA; while Kaforou_27 yielded the highest weighted AUCs of (0.93, 95% CI: 0.87 - 0.97) using PLAGE (Table S3).

Overall, 67 gene signatures demonstrated weighted AUCs ≥ 0.80 in at least one comparison using either ssGSEA or PLAGE (Figure 3A). Notably, 43 (64.2%) gene signatures had weighted AUCs ≥ 0.80 in distinguishing patients with PTB from either Control or LTBI, irrespective of the evaluation methods (PLAGE or ssGSEA). Additionally, eight (11.9%) gene signatures with weighted AUCs ≥ 0.80 were found in both comparisons using PLAGE (Figure 3A). Moreover, 35 gene signatures had weighted AUCs ≥ 0.90 in at least one comparison using either ssGSEA or PLAGE, with four (11.4%) gene signatures (Blankley_5, Kaforou_27, Kaul_3, and Tabone_RES_27) consistently maintaining weighted AUCs ≥ 0.90 across all four comparisons (Figure 3B). In summary, a greater number of gene signatures showed high predictive ability (weighted AUCs ≥ 0.90) in PTB vs. Control (n = 33) compared to PTB vs. LTBI (n = 22; Figure 3B).

**Figure 3.**
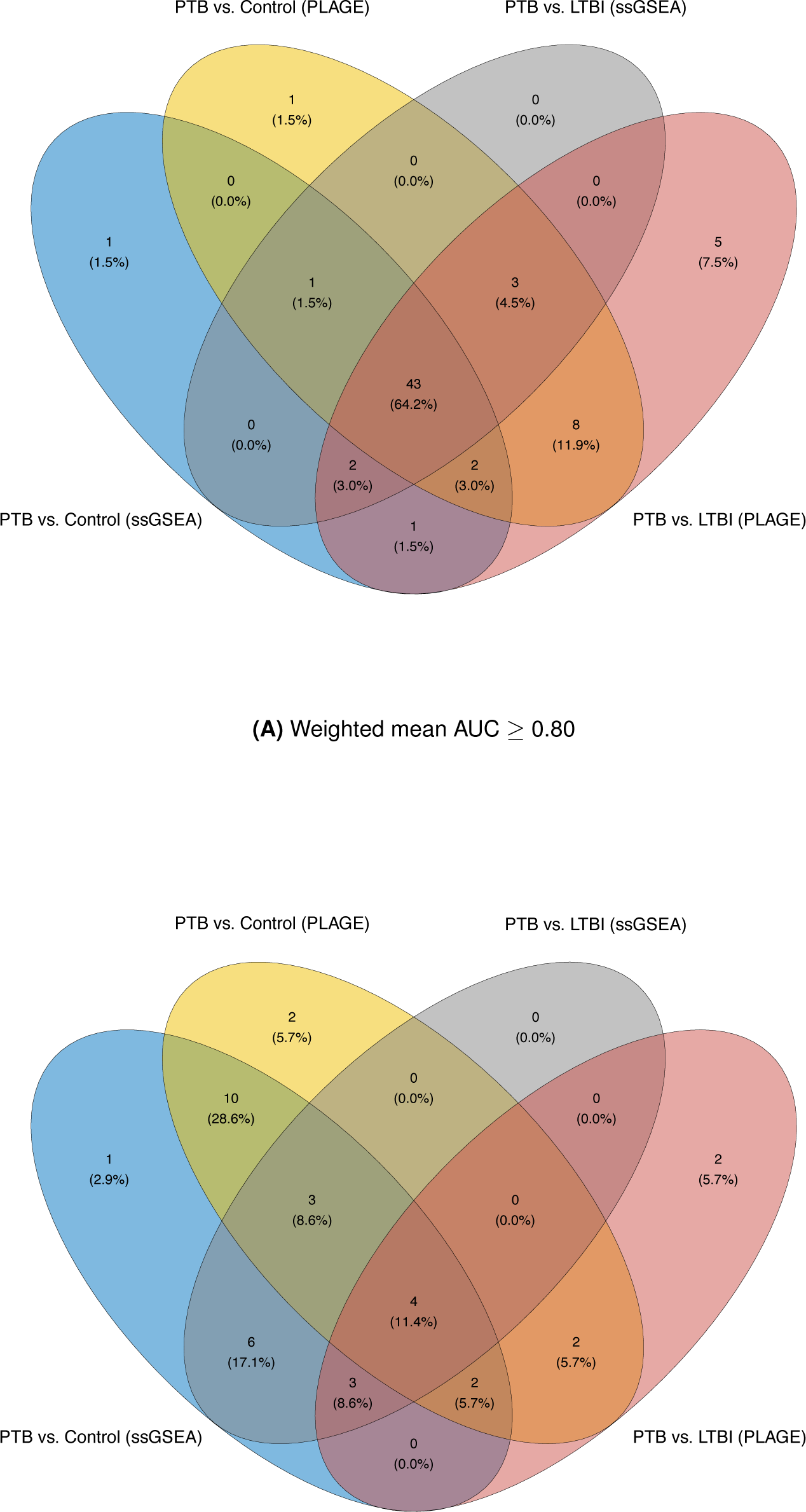
The number and percentage of overlapping TB gene signatures with weighted mean AUC value *≥* 0.80 **(A)** and weighted AUC value *≥* 0.90 **(B)** for comparisons of PTB versus controls and PTB versus LTBI using ssGSEA and PLAGE.

For PTB vs. Control, the highest weighted sensitivity was observed with Kaul_3 (0.92, 95% CI: 0.80 - 0.99) using ssGSEA, while Maertzdorf_15 and PennNich_RISK_6 achieved a weighted sensitivity value of 0.91 (95% CI: 0.75 - 0.99) using PLAGE (Table S4). Additionally, Blankley_5, Hoang_OD_20, and Roe_3 demonstrated the highest weighted specificity of 0.94 (95% CI: 0.81 - 1.00) using ssGSEA, and the highest weighted specificity was given by Natarajan_7 (0.94, 95% CI: 0.83 - 0.99) using PLAGE (Table S5).

For PTB vs. LTBI, Thompson_9 exhibited the highest weighted sensitivity (0.87, 95% CI: 0.71 - 0.97) when evaluated by ssGSEA, and Suliman_4 had the highest weighted sensitivity (0.88, 95% CI: 0.74 - 0.98) using PLAGE (Table S4). Moreover, the highest weighted specificity was given by Bloom_OD_144 and Tornheim_71, both recording a value of 0.92 (95% CI: 0.77 - 0.99) using ssGSEA, and Thompson_9 achieved the highest weighted specificity of 0.94 (95% CI: 0.81 - 1.00) using PLAGE (Table S5).

### Individual Performance of TB Gene Signatures

The evaluation of gene signature performance within individual datasets is presented in Figure S1 and Figure S2. In the context of PTB vs. Control, both ssGSEA and PLAGE encountered difficulties computing profiling scores for Kulkarni_HIV_2 (*INSL3, RAB20*) in three out of 25 datasets. Notably, either one of the genes were not found in GSE6112 and Bruno study (31), while both genes were not identified in GSE74092 (Figure S1). For PTB vs. Control, the predictive performance of seven gene signatures were unavailable in GSE6112, mainly because the use of the MPIIB custom human sequencing platform. Additionally, profiling socres for 13 gene signatures profiling score were not accessible in GSE74092, given its utilization of an RT-PCR assay (Figure S1). Regarding the comparison of PTB vs. LTBI, results for six gene signatures (Tabone_OD_11, Gong_OD_20, Kulkarni_HIV_2, Suliman_4, Roe_3, and Kaul_3) were unavailable in GSE6112 (Figure S2).

### Ensemble of multiple gene signatures

Ensembles of multiple gene signatures in differentiating subjects with PTB from Control or from LTBI were also implemented as an application and compared using the curatedTBData package. In this practice, 30 ensemble sets with each set contain 5 gene signatures had been evaluated for both comparisons, and as expected, the performance of ensemble gene signatures varied greatly set by set (Figure S3; Figure S6). Overall, for the comparison of PTB vs. Control, 25 out of 30 ensemble sets demonstrated higher weighted AUCs when compared to their single best gene signature, whether evaluated using PLAGE (n = 17) or ssGSEA (n = 18) (Figure S3A). Specifically, 8 out of 17 sets had smaller weighted SD using PLAGE, and 17 out of 18 sets had smaller weighted SD using ssGSEA (Figure S6A). In the context of PTB vs. LTBI, all ensemble sets outperformed their single best subset, whether assessed through PLAGE (n = 6) or ssGSEA (n = 29; Figure S3B). Among these sets, 5 out 6 had lower value of weighted SD with PLAGE, and 14 out 16 had lower value of weighted SD with ssGSEA (Figure S6B). Three selected ensemble sets were highlighted to illustrate the distinct performance of ensembling biomarkers (Figure 4, 5).

**Figure 4.**
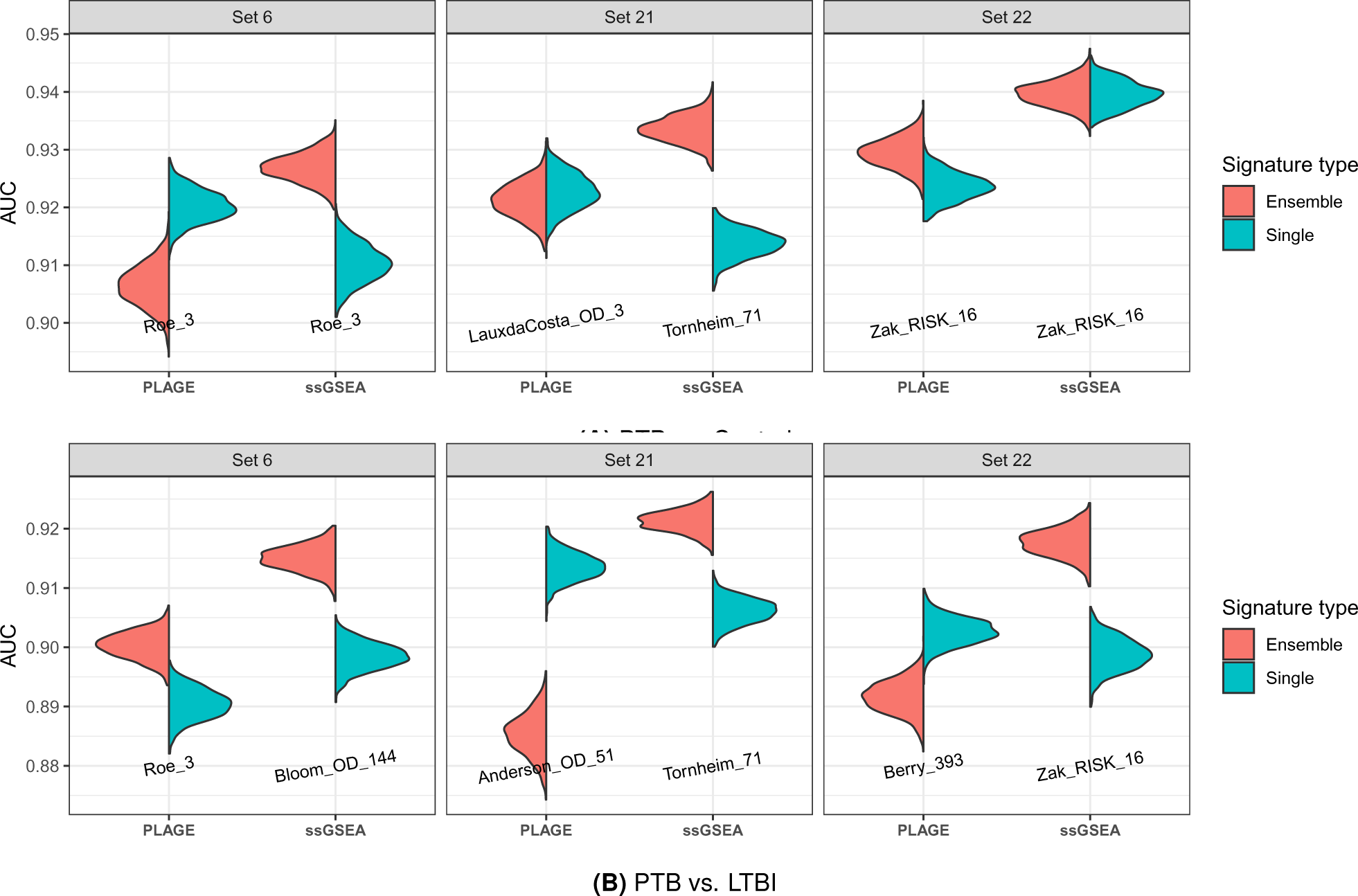
Comparison of selected ensemble of multiple signatures to its the single best signature in distinguishing active TB from healthy control **(A)** and from LTBI **(B)** using PLAGE or ssGSEA. Each set contains 5 TB gene signatures that were randomly selected from signature list. AUC distribution were computed based on 300 cross-study validation. The name of the gene signature that had the highest weighted mean AUC values within the corresponding set was labeled on the plot.

### Active TB vs. Control

Ensemble *Set 6* exhibited a weighted AUC of 0.90 (95% CI: 0.89 - 0.91) when evaluated by PLAGE, a value significantly smaller than its single best gene signature, Roe_3 (0.92, 95% CI: 0.91 - 0.93; p-value = 9.0*e* − 3). Additionally, *Set 6* demonstrated a smaller value of weighted SD (*SD_w_* = 0.024) in comparison to Roe_3 (*SD_w_* = 0.021) and Anderson_42 (*SD_w_* = 0.021) using PLAGE. When assessed by ssGSEA, this *Set 6* had significantly higher weighted AUCs (0.93, 95% CI: 0.92 - 0.94) when compared to its best subset Roe_3 (0.91, 95% CI: 0.90 - 0.92; p-value *<* 1.0*e* − 3; Figure 4A), and had the smallest weighted SD (*SD_w_* = 0.016) when evaluated by ssGSEA (Figure 5A).

**Figure 5.**
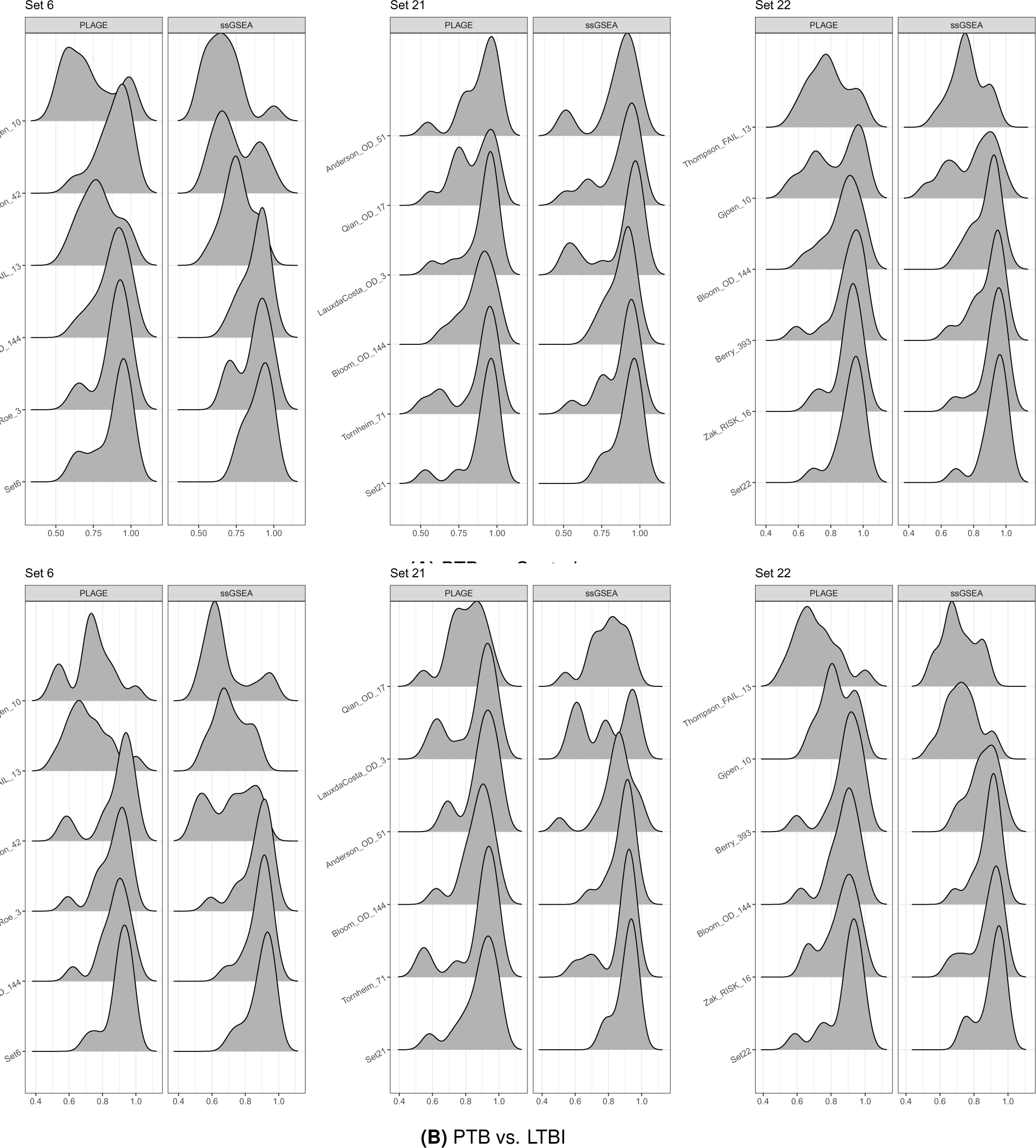
Distribution of AUC values for selected ensemble gene sets and gene signatures within the sets across evaluated datasets for PTB vs. Control **(A)** and PTB vs. LTBI **(B)**.

Although *Set 21* displayed similar weighted AUCs to its highest subset, LauxdaCosta_OD_3 (0.92, 95% CI: 0.91 - 0.93; p-value = 0.78) according to PLAGE (Figure 4A), it had the smallest weighted SD (*SD_w_* = 0.024) when compared to all other gene signatures within the ensemble set (Figure 5A). When evaluated by ssGSEA, *Set 21* significantly outperformed its single best gene signature, Tornheim_71 (0.91, 95% CI: 0.90 - 0.93; p-value *<* 1.0*e* −3; Figure 4A) and as anticipated had the lowest weighted SD among all the five gene sets (*SD_w_* = 0.016; Figure 5A). For *Set 22*, its weighted AUCs were comparable to the results from Zak_RISK_16 using either PLAGE or ssGSEA (p-value *>* 0.05 for both methods; Figure 4A). However, *Set 22* had the smallest weighted SD when compared to any of gene signature within the ensemble set for both methods (*SD_w_* = 0.016 for PLAGE and *SD_w_* = 0.015 for ssGSEA; Figure 5A).

### Active TB vs. LTBI

Ensemble *Set 6*, characterized by a weighted SD of 0.014 for PLAGE and 0.015 for ssGSEA, demonstrated superior performance compared to its single best gene signature, Roe_3 (*SD_w_* = 0.019) and Bloom_OD_144 (*SD_w_* = 0.016) using both PLGAE and ssGSEA (p-value *<* 0.01 for both methods; Figure 4B; Figure S6B). The performance of *Set 21* (0.88, 95% CI: 0.87 - 0.88, *SD_w_* = 0.022) was significantly inferior to its best subset Anderson_OD_51 (0.91, 95% CI: 0.90 - 0.93, *SD_w_* = 0.019) using PLAGE (p-value *<* 1.0*e* − 3; Figure 5B). Conversely, when ssGSEA was the profiling method, *Set 21* (0.92, 95% CI: 0.91 - 0.93) significantly outperformed its best subset, Tornheim_71 (p-value *<* 1.0*e* − 3, *SD_w_* = 0.019; Figure 4B), with the smallest weighted SD of 0.012 (Figure 5B). Lastly, *Set 22* (*SD_w_* = 0.021) exhibited smaller weighted AUCs compared to its best subset, Berry_393 using PLAGE (p-value = 0.011, *SD_w_* = 0.019; Figure 4B, 5B). In contrast, *Set 22* (*SD_w_* = 0.016) outperformed it best subset, Zak_RISK_16 (*SD_w_* = 0.020) using ssGSEA, with the smallest weighted SD among all of its subsets (p-value *<* 1.0*e* − 3; Figure 4B, 5B).

## Discussion

In this paper, we introduced the curatedTBData data package for R statistical programming environment. Currently, this package includes 49 TB transcriptomic datasets with well-annotated metadata, which serves as a curated resource that allows researchers to make efficient use of currently existing TB gene expression profiles. One of the most important outstanding problems for public transcriptomic TB datasets is that some of studies lack clinical information, and others may possess inconsistent definition of clinical features across different datasets (32). In our curation efforts, we were able to retrieve important clinical characteristics including age, gender, geographical region, TB status, etc., for most of the studies. However, features such as TB diagnostic methods remained inconsistent across many of the studies due to their original design. Currently, there are various techniques to diagnose TB outcomes including but not limited to sputum culture, sputum PCR, sputum microscopy, bronchoalveolar lavage (bal) culture, and biopsies. LTBI is assessed using tuberculin skin test (TST), and Interferon-Gamma Release Assays (IGRAs), which – if positive – reflect exposure, but are unable to speak to past infection that has resolved as opposed to ongoing latent infection. Most datasets used either one or combined technique(s) to identify a patient’s TB status, while there are some studies that do not contain TB diagnostic information (Table S2). The relatively large number of well-annotated studies in this database could contribute to future work that recover missing clinical annotations.

While there are several approaches that discussed the problems about dealing with multiplicative Affymetrix probesets or Ensembl IDs (33); in the current version of the data package, we used the naive method by taking the median value of the probes/Ensembl IDs that target the same gene. For the output of the MultiassayExperiment object, the updated gene expression data with gene symbols as row names were showed in the slot with name assay_curated, and the un-collapsed version of the data were also provided in the slot with assay_raw. In summary, this heterogeneous database is a representation of clinical practice as patients were diagnosed using different methods, and it also represents heterogeneity in both host and pathogen genetics (2).

In this study, we compared 72 TB gene signatures that distinguish subjects with PTB from LTBI or PTB from healthy controls using the curatedTBData database. Our analysis illustrated that larger number of TB gene signatures had high diagnostic ability (weighted AUCs ≥ 0.90) in the comparison of PTB vs. Control when compared to PTB vs. LTBI (Table S3; Figure 3). Meanwhile, the selection of profiling methods did not have significant impact on gene signatures’ predictive ability in PTB vs. Control (p-value = 0.12). In contrast, the results given by PLAGE outperformed its counterpart from ssGSEA in PTB vs. LTBI (p-value ≤ 1.0*e* −3; Table S3; Figure 3). One explanation for such outperformance was that gene signatures were pre-selected based on prior knowledge, and using dimension reduction method like SVD on the activity levels of the gene signatures could reveal their biological meaning (25). On the contrary, it has been shown that ssGSEA was likely to give null results when both upregulated and downregulated subsets of genes were presented within the gene sets (34).

Generally, ensemble learning of multiple TB gene signatures were able to improve the overall predictive ability. This outperformance was expected, as ensemble stacking technique combines information from each of single learner, yielding higher accuracy results (35). Additionally, the performance of ensemble sets in PTB vs. LTBI exhibited superiority compared to the PTB vs. Control (Figure S3). It is noteworthy that the results were not directly comparable due to different samples were included in theses two comparisons. From an ensemble stacking perspective, gene signatures in the context of PTB vs. LTBI may be considered as weak learners in contrast to PTB vs. Control (Figure 3), and the advantage of ensemble technique lies in enhancing model performance by combining weak learners (35; 36).

In our practice, large weights were assigned to those with high predictive ability on average (Table S3; Figure S4). For example, gene signatures like Zak_RISK_16, Kaforou_27, and Blankley_5 etc., had consistently non-zero weights whenever they were included in the ensemble sets(Figure S4). One large deficient were shown in *set 28* with PLAGE (Figure S3) for both comparisons. This was primarily because *set 28* mistakenly shrank the coefficient for signatures with high predictive ability to zero (i.e. Berry_393 in PTB vs. Control and Estevez_133 in PTB vs. LTBI) (Figure S3), which also suggested the instability of using PLAGE in the context of stacking ensemble.

Although the outperformance of ensemble sets was trivial in terms of weighted AUCs (outperformance up to 3%), the consistency of the results were increased by leveraging ensembling strategy (Figure S6). Except for PTB vs. Control using PLAGE, where only 8 out of 17 sets had lower value of weighted SD, all the rest of comparisons yielded high percentage (*>* 83%) of ensemble sets with increased stability when compared to their corresponding single best subset (Figure S6). The reduction of variation for model is another critical pattern of ensemble strategy. It has also been shown in some empirical studies that the improvement of model performance was attributed to the reduction of variance to some extents in voting algorithm including Bagging and Boosting (36; 37).

For both comparisons, the performance of ensemble sets given by ssGSEA was better than the performance given by PLAGE. The outperformance of ensemble of gene signatures using ssGSEA was possibly due to the results scores given by ssGSEA were rank-based and single-sample, which were robust against heterogeneous datasets and does not depend on other samples in the datasets (34). Moreover, large variations of the profiling scores were given by ssGSEA (*SD_P T B_* = 0.17*, SD_LT BI_* = 0.154*, SD_Control_* = 0.15) when compared to PLAGE (*SD_P T B_* = 0.033*, SD_LT BI_* = 0.034*, SD_Control_* = 0.031; Figure S5), which is critical for the outperformance of ensemble learning (38). In addition, the coefficient for selected gene signatures were more likely to be positive using ssGSEA when compared to the results from PLAGE (Figure S4), this is largely due the resulting scores given by ssGSEA for most of gene signatures in ensemble, on average, were higher for PTB when compared to Control or LTBI (except Anderson_42 and Verhagen_10 for PTB vs. Control, and Suliman_RISK_4 and Thompson_FAIL_13 for both comparisons; Figure S5).

## Conclusion

### Contribution Summary

The curatedTBData package offers an extensive resource of curated gene expression and clinically annotated data for various applications, including but not limited to the comparative analysis of existing TB gene signatures and the discovery of new TB gene signatures. This database provides researchers to make efficient use of TB gene expression data and offers a resource for reproducible and replicable analysis of the TB transcriptomic datasets.

In the context of TB diagnostics using blood-based biomarkers, it is important to evaluate existing gene signatures in a systematic way. Our comparative analysis showed that 65 out of 72 gene signatures were able to separate patients with PTB from healthy controls or LTBI with high accuracy (weighted AUCs ≥ 0.8) using either one of the two gene set scoring methods, and gene signatures tend to have higher accuracy in distinguishing PTB from Control when comparing to PTB from LTBI (Figure 3). Our results provided strong evidence for the diagnosis of TB using blood-based biomarkers, as well as the evaluation of TB gene signatures using gene set scoring methods. We also showed that predictive ability can be improved or provide a better lower bound on performance by leveraging stacking ensemble strategy. This technique can be highly recommended for TB diagnosis using blood-based biomarker, especially for heterogeneous testing cohort that come from different sources.

### Limitation and Next Steps

Our analysis has a few limitations. When building curatedTBData, the mapping of probe sets to common gene symbols is critical for promoting cross-study consistency. However, gene identification is a persistent problem in integration analysis (10). We primarily used information from GEO as the reference, but that information may have updated with the development of sequencing technology. Also, when evaluating the performance of 72 gene signatures, we did not use the discovery model for each signature from its original study. For some gene signatures, certain underlying information may be lost if we used gene set enrichment analysis to evaluate their performance. Moreover, for some gene signatures, some genes were not available across different studies. Although we used the R package HGNCHelper (27) to update both gene signatures and gene expression profile ID, this may reduce their ability to distinguish TB subtypes across multiple studies.

## Online data supplement

**Table S1:** Transcriptome datasets in the curatedTBData package.

| Study | Platform | Geographical Region | Age | Diagnosis Method | Control | LTBI | PTB | OD | Total |
| --- | --- | --- | --- | --- | --- | --- | --- | --- | --- |
| GSE31348 [1] | GPL570 | South Africa | 18-65 | NA | NA | NA | 27 | NA | 27 |
| GSE36238 [1] | GPL570 | South Africa | 18-65 | Sputum culture | NA | 9 | NA | NS | 9 |
| GSE41055 [2] | GPL5175 | Venezuela | 1-15 | Sputum culture, TST, IGRA(QFT-GIT) | 9 | 9 | 9 | NA | 27 |
| GSE54992 [3] | GPL570 | China | 18-68 | Sputum culture, chest radiography | 6 | 6 | 9 | NA | 21 |
| GSE73408 [4] | GPL11532 | US | >18 | NA | NA | 35 | 35 | 39 | 109 |
| GSE107731 | GPL15207 | Mongolian | NA | NA | 3 | NA | 3 | NA | 6 |
| GSE79362 [5] | GPL11154 | South Africa | 12-18 | Sputum culture, Sputum Smears | NA | 104 | 40 | NA | 144 |
| GSE84076 [6] | GPL16791 | Brazil | >18 | Sputum culture, Sputum microscopy | 12 | 16 | 8 | NA | 36 |
| GSE89403 | GPL11154 | South Africa | NA | Sputum culture, MGIT, XPERT, TGRV | 21 | NA | 100 | 17 | 138 |
| GSE94438 [7] | GPL11153 | South Africa, The Gambia, Ethiopia, Uganda | 10-18 | Sputum culture, Smear microscopy, Clinical signs | NA | 259 | 75 | NA | 334 |
| GSE107991 [8] | GPL20301 | UK | >18 | Sputum culture | NA | 33 | 21 | NA | 54 |
| GSE107992 [8] | GPL20301 | South Africa | >18 | Sputum culture | NA | 31 | 16 | NA | 47 |
| GSE107993 [8] | GPL20301 | UK | 17-65 | Sputum culture | 15 | 16 | NA | NA | 31 |
| GSE107994 [8] | GPL20301 | UK | 16-84 | Sputum culture | 50 | 58 | 53 | NA | 161 |
| GSE101705 [9, 10, 11] | GPL18573 | South India | >18<sup>a</sup> | sputum AFB and culture, TST | NA | 16 | 28 | NA | 44 |
| GSE107104<sup>b</sup> [12] | GPL11154 | NA | >18 | Smear or microbiologically-confirmed TB | 16 | NA | 17 | NA | 33 |
| GSE112104 [13] | GPL16791 | Brazil | 1-72 | Sputum culture | NA | 21 | 30 | NA | 51 |
| Tornheim<sup>c</sup> [14] | GPL16791 | India | 2-14 | TST, IGRA | 32 | NA | 8 | NA | 40 |
| GSE19435 [15] | GPL6947 | NA | 20-55 | Sputum culture etc. | 12 | NA | 7 | NA | 19 |
| GSE19439 [15] | GPL6947 | UK | 19-72 | Sputum culture etc. | 12 | 17 | 13 | NA | 42 |
| GSE19442 [15] | GPL6947 | South Africa | 18-48 | Sputum culture etc. | NA | 31 | 20 | NA | 51 |
| GSE19443 [15] | GPL6947 | UK | >18 | Sputum culture etc. | 4 | NA | 7 | NA | 11 |
| GSE19444 [15] | GPL6947 | UK | >18 | Sputum culture etc. | 12 | 21 | 21 | NA | 54 |
| GSE22098 [15] | GPL6947 | NA | 1-84 | Sputum culture etc. | 81 | NA | NA | 193 | 274 |
| GSE29536 [16, 17, 18] | GPL6102 | UK | NA | Sputum culture, chest radiography | 6 | NA | 9 | NA | 15 |
| GSE37250<sup>b</sup> [19] | GPL10558 | South Africa, Malawi | >17 | Sputum culture, chest radiography | NA | 167 | 195 | 175 | 537 |
| GSE39939<sup>b</sup> [20] | GPL10558 | Kenya | <15 | Sputum culture, chest radiography, TST | NA | 14 | 79 | 64 | 157 |
| GSE39940<sup>b</sup> [20] | GPL10558 | South Africa, Malawi | >15 | Sputum culture, chest radiography, TST | NA | 54 | 111 | 169 | 334 |

Table S1 – Continued from previous page
| Study | Platform | Geographical Region | Age | Diagnosis Method | Control | LTBI | PTB | OD | Total |
| --- | --- | --- | --- | --- | --- | --- | --- | --- | --- |
| GSE40553<br>[21] | GPL10558 | South Africa, UK | NA | NA | NA | 38 | 37 | NA | 75 |
| GSE42825<br>[22] | GPL10558 | Germany | >17 | Sputum culture | 23 | NA | 8 | 11 | 42 |
| GSE42826<br>[22] | GPL10558 | Germany | >17 | Sputum culture | 52 | NA | 11 | 39 | 102 |
| GSE42827<br>[22] | GPL10558 | Germany | >17 | NA | NA | NA | NA | 5 | 5 |
| GSE42830<br>[22] | GPL10558 | Germany | >17 | Sputum culture | 38 | NA | 16 | 41 | 95 |
| GSE42831<br>[22] | GPL10558 | Germany | >17 | NA | NA | NA | NA | 7 | 7 |
| GSE42832<br>[22] | GPL10558 | Germany | >17 | Sputum culture | 5 | NA | 5 | 5 | 15 |
| GSE50834<sup>b</sup><br>[23] | GPL10558 | South Africa | 30-40 | NA | 22 | 21 | NA | NA | 43 |
| GSE56153<br>[24] | GPL6883 | Indonesia | >15 | Sputum microscopy, chest X-ray | 23 | NA | 23 | NA | 46 |
| GSE69581<sup>b</sup><br>[25] | GPL10558 | South Africa | >17 | Sputum culture, sputum Xpert MTB-RIF | NA | 25 | 15 | NA | 40 |
| GSE83456<br>[26] | GPL10558 | UK | NA | Sputum culture etc. | 61 | NA | 45 | 49 | 155 |
| GSE83892<sup>b</sup><br>[27] | GPL10558 | UK | >17 | NA | 17 | NA | 33 | NA | 50 |
| Bruno<sup>c</sup> [28] | GPL10558 | India | 25-60 | Sputum culture | 29 | NA | 53 | 28 | 110 |
| GSE25534 | GPL1708 | NA | NA | Sputum culture, chest radiograph, TST | 13 | 45 | 44 | NA | 102 |
| GSE28623<br>[29, 30] | GPL4133 | The Gambia | 16-53 | Sputum microscopy, chest X-ray | 37 | 25 | 46 | NA | 108 |
| GSE34608<br>[31] | GPL6480 | Germany | 17-73 | NA | 18 | NA | 8 | 18 | 44 |
| GSE62147<br>[32] | GPL6480 | The Gambia | NA | Sputum culture, sputum smears | NA | NA | 14 | 12 | 26 |
| GSE81746 | GPL17077 | India | 25-65 | NA | 2 | 1 | 5 | NA | 8 |
| GSE62525<br>[33] | GPL19651 | Taiwan | NA | NA | 7 | 7 | 7 | NA | 21 |
| GSE74092<br>[34] | GPL21040 | India | NA | GenXpert assay | 76 | NA | 113 | NA | 189 |
| GSE6112<br>[35] | GPL4475 | NA | NA | Sputum culture, chest X-ray | 36 | 17 | 19 | NA | 72 |

<sup>a</sup> One subject is 16 years old<sup>b</sup> Contain HIV positive subjects<sup>c</sup> Gene Expression Omnibus Identifier not available

**Table S2:**
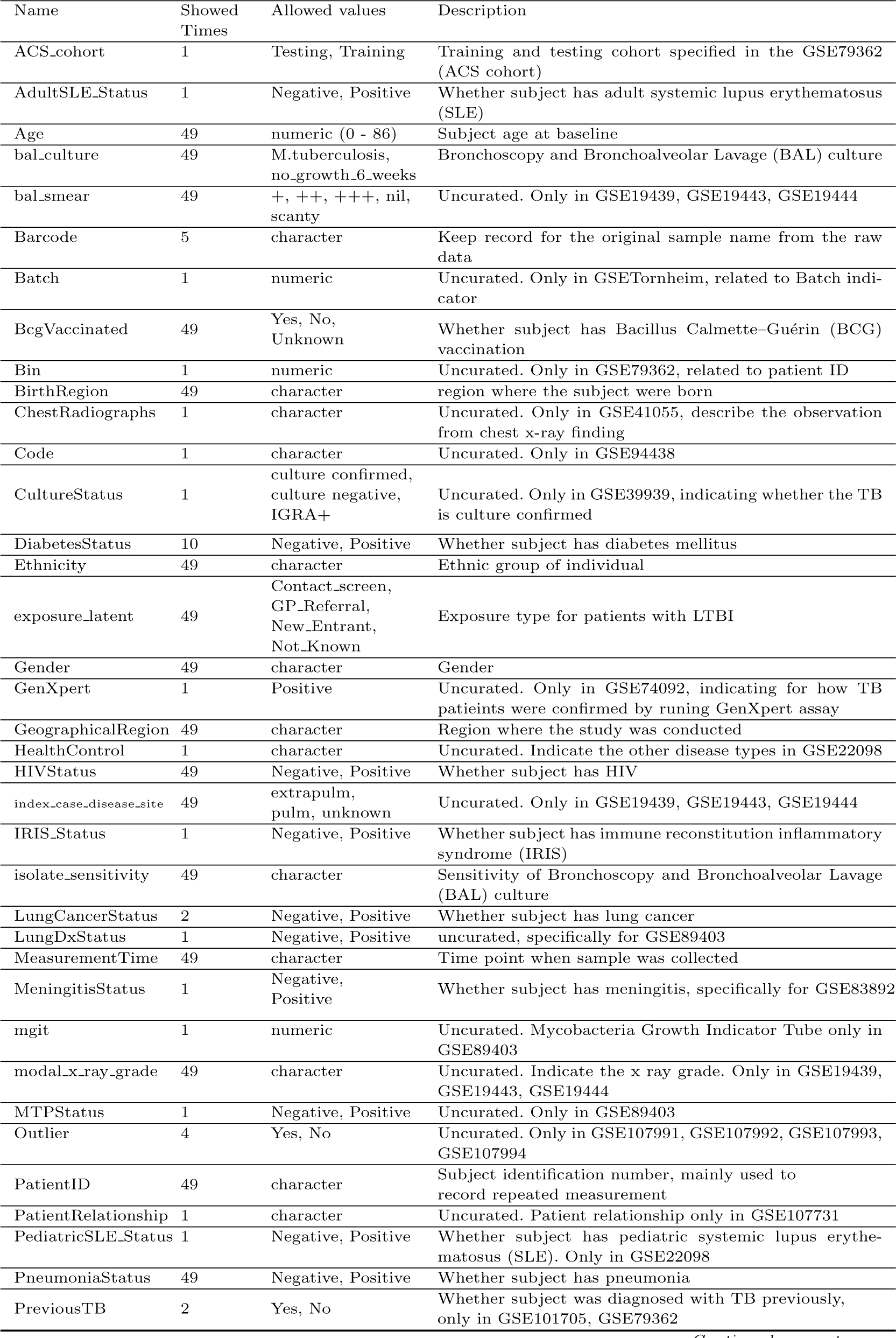
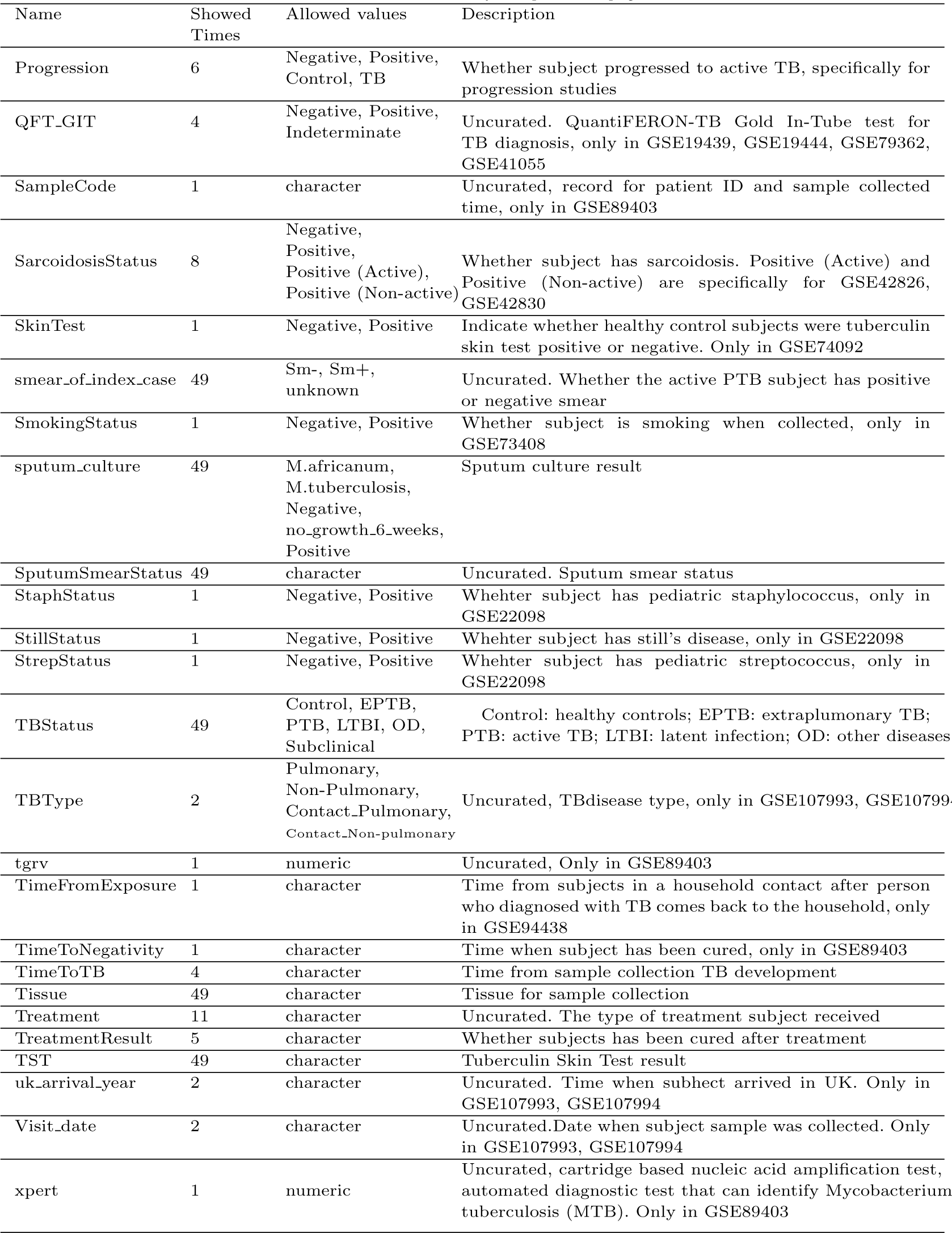
Description of clinical annotations.

**Table S3:**
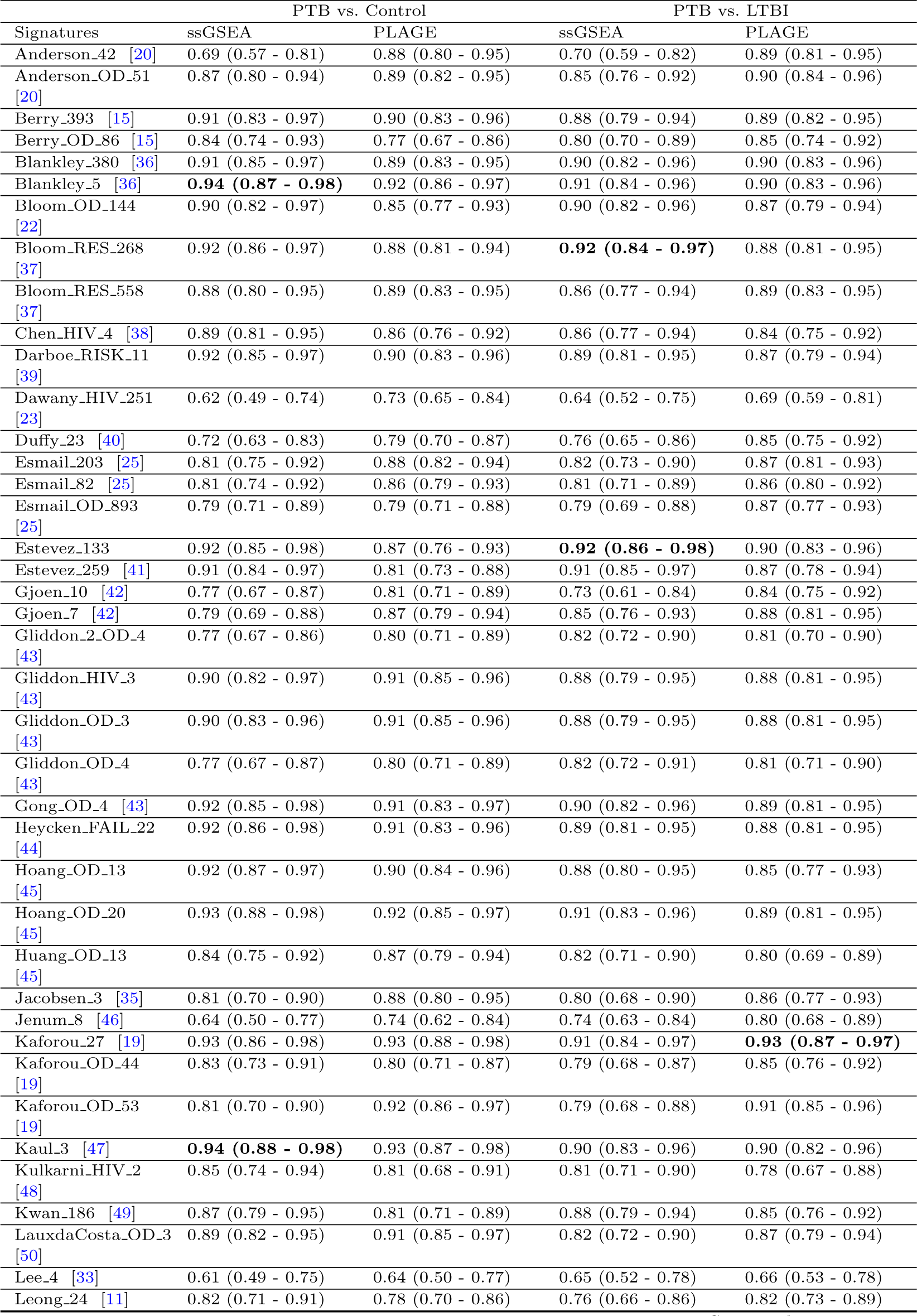
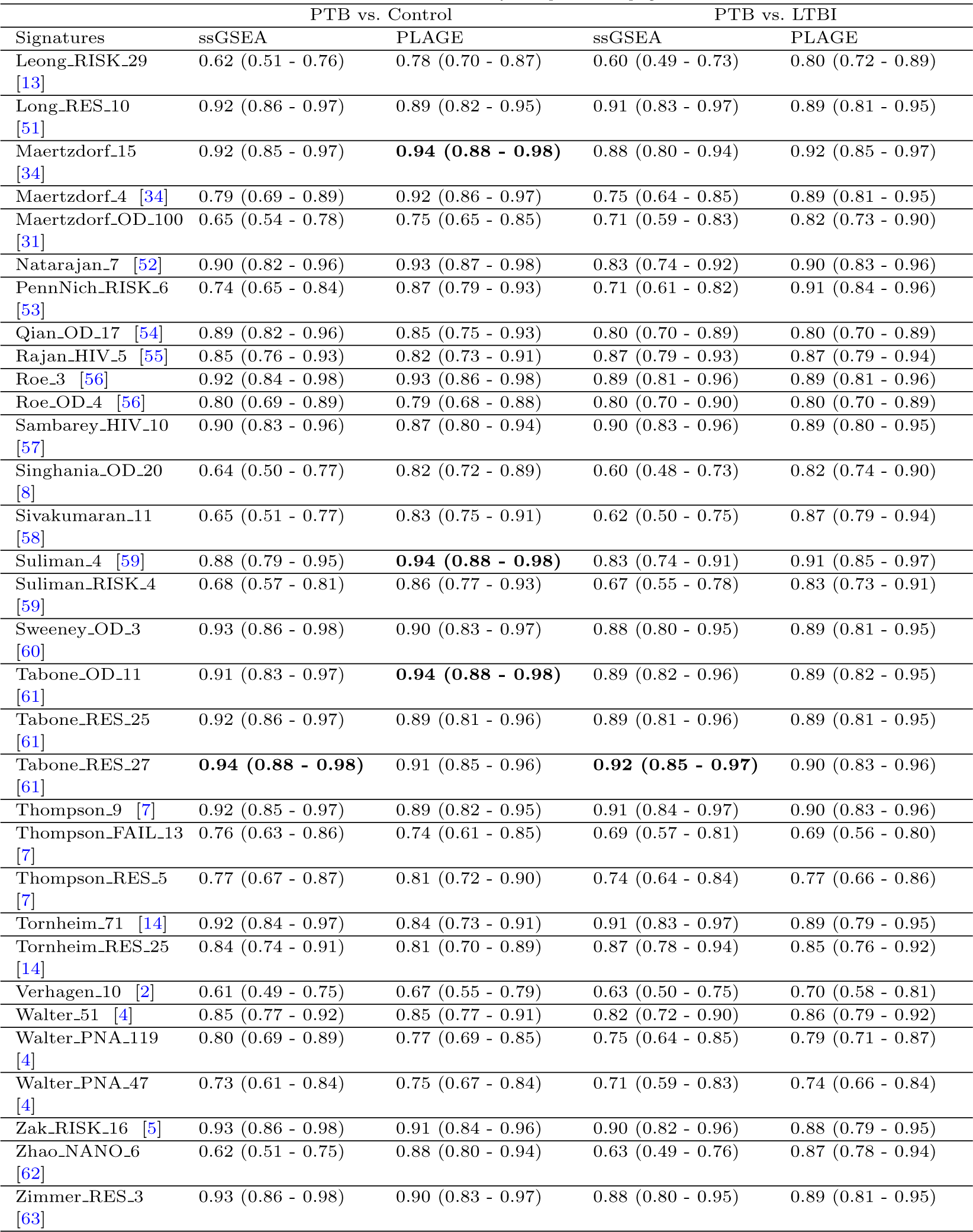
Weighted mean AUC and 95% confidence interval for 72 signatures for distinguishing patients with active TB from healthy controls or from LTBI.

**Table S4:** Weighted mean Sensitivity and 95% confidence interval for 72 signatures for distinguishing patients with active TB from healthy controls or from LTBI.

| Signatures | PTB vs. Control |  | PTB vs. LTBI |  |
| --- | --- | --- | --- | --- |
|  | ssGSEA | PLAGE | ssGSEA | PLAGE |
| Anderson_42 | 0.68 (0.39 - 0.90) | 0.86 (0.68 - 0.98) | 0.64 (0.45 - 0.93) | 0.84 (0.69 - 0.97) |
| Anderson_OD_51 | 0.84 (0.68 - 0.97) | 0.82 (0.69 - 0.96) | 0.75 (0.58 - 0.92) | 0.84 (0.68 - 0.95) |
| Berry_393 | 0.85 (0.71 - 0.96) | 0.86 (0.74 - 0.97) | 0.79 (0.64 - 0.95) | 0.80 (0.67 - 0.95) |
| Berry_OD_86 | 0.79 (0.63 - 0.95) | 0.70 (0.58 - 0.92) | 0.75 (0.53 - 0.93) | 0.81 (0.62 - 0.96) |
| Blankley_380 | 0.89 (0.73 - 0.98) | 0.83 (0.71 - 0.96) | 0.80 (0.67 - 0.95) | 0.82 (0.68 - 0.95) |
| Blankley_5 | 0.87 (0.76 - 0.98) | 0.86 (0.75 - 0.97) | 0.84 (0.69 - 0.96) | 0.83 (0.69 - 0.97) |
| Bloom_OD_144 | 0.81 (0.69 - 0.97) | 0.83 (0.64 - 0.94) | 0.82 (0.68 - 0.95) | 0.79 (0.61 - 0.94) |
| Bloom_RES_268 | 0.86 (0.75 - 0.97) | 0.81 (0.70 - 0.95) | 0.83 (0.69 - 0.96) | 0.80 (0.65 - 0.94) |
| Bloom_RES_558 | 0.82 (0.66 - 0.94) | 0.85 (0.73 - 0.96) | 0.77 (0.61 - 0.94) | 0.81 (0.68 - 0.95) |
| Chen_HIV_4 | 0.85 (0.74 - 0.95) | 0.85 (0.71 - 0.97) | 0.83 (0.66 - 0.96) | 0.78 (0.63 - 0.94) |
| Darboe_RISK_11 | 0.87 (0.75 - 0.98) | 0.87 (0.73 - 0.98) | 0.82 (0.67 - 0.95) | 0.85 (0.66 - 0.95) |
| Dawany_HIV_251 | 0.67 (0.33 - 0.91) | 0.76 (0.55 - 0.94) | 0.64 (0.34 - 0.90) | 0.65 (0.46 - 0.92) |
| Duffy_23 | 0.63 (0.45 - 0.87) | 0.77 (0.61 - 0.93) | 0.68 (0.50 - 0.90) | 0.81 (0.63 - 0.97) |
| Esmail_203 | 0.77 (0.65 - 0.93) | 0.86 (0.70 - 0.97) | 0.78 (0.57 - 0.93) | 0.85 (0.66 - 0.98) |
| Esmail_82 | 0.79 (0.63 - 0.97) | 0.82 (0.69 - 0.97) | 0.74 (0.57 - 0.93) | 0.79 (0.64 - 0.97) |
| Esmail_OD_893 | 0.74 (0.55 - 0.91) | 0.78 (0.61 - 0.94) | 0.69 (0.50 - 0.89) | 0.83 (0.65 - 0.96) |
| Estevez_133 | 0.88 (0.74 - 0.98) | 0.80 (0.66 - 0.91) | 0.86 (0.72 - 0.98) | 0.81 (0.67 - 0.98) |
| Estevez_259 | 0.84 (0.72 - 0.97) | 0.74 (0.63 - 0.95) | 0.84 (0.69 - 0.97) | 0.83 (0.65 - 0.98) |
| Gjoen_10 | 0.72 (0.52 - 0.90) | 0.74 (0.55 - 0.92) | 0.67 (0.46 - 0.93) | 0.77 (0.60 - 0.95) |
| Gjoen_7 | 0.69 (0.54 - 0.89) | 0.82 (0.67 - 0.94) | 0.81 (0.59 - 0.95) | 0.84 (0.67 - 0.96) |
| Gliddon_2_OD_4 | 0.71 (0.52 - 0.90) | 0.69 (0.53 - 0.91) | 0.78 (0.59 - 0.95) | 0.79 (0.58 - 0.96) |
| Gliddon_HIV_3 | 0.84 (0.72 - 0.96) | 0.87 (0.74 - 0.96) | 0.82 (0.67 - 0.96) | 0.80 (0.64 - 0.96) |
| Gliddon_OD_3 | 0.84 (0.71 - 0.97) | 0.87 (0.74 - 0.96) | 0.82 (0.67 - 0.96) | 0.80 (0.64 - 0.96) |
| Gliddon_OD_4 | 0.71 (0.51 - 0.90) | 0.69 (0.53 - 0.91) | 0.78 (0.59 - 0.95) | 0.79 (0.58 - 0.96) |
| Gong_OD_4 | 0.87 (0.76 - 0.97) | 0.87 (0.73 - 0.97) | 0.84 (0.68 - 0.96) | 0.82 (0.67 - 0.96) |
| Heycken_FAIL_22 | 0.87 (0.75 - 0.98) | 0.83 (0.72 - 0.96) | 0.80 (0.67 - 0.96) | 0.80 (0.63 - 0.95) |
| Hoang_OD_13 | 0.86 (0.74 - 0.97) | 0.87 (0.75 - 0.98) | 0.82 (0.66 - 0.96) | 0.82 (0.66 - 0.95) |
| Hoang_OD_20 | 0.88 (0.79 - 0.98) | 0.87 (0.74 - 0.99) | 0.86 (0.68 - 0.97) | 0.82 (0.67 - 0.96) |
| Huang_OD_13 | 0.78 (0.60 - 0.95) | 0.84 (0.67 - 0.97) | 0.80 (0.56 - 0.95) | 0.69 (0.54 - 0.95) |
| Jacobsen_3 | 0.84 (0.60 - 0.95) | 0.86 (0.71 - 0.96) | 0.78 (0.61 - 0.96) | 0.80 (0.64 - 0.94) |
| Jenum_8 | 0.68 (0.32 - 0.95) | 0.73 (0.53 - 0.89) | 0.74 (0.48 - 0.91) | 0.75 (0.53 - 0.92) |
| Kaforou_27 | 0.88 (0.77 - 0.98) | 0.88 (0.79 - 0.98) | 0.86 (0.73 - 0.98) | 0.86 (0.72 - 0.98) |
| Kaforou_OD_44 | 0.76 (0.63 - 0.95) | 0.76 (0.64 - 0.91) | 0.71 (0.53 - 0.89) | 0.83 (0.65 - 0.96) |
| Kaforou_OD_53 | 0.72 (0.56 - 0.92) | 0.89 (0.78 - 0.97) | 0.71 (0.51 - 0.93) | 0.85 (0.72 - 0.97) |
| Kaul_3 | 0.92 (0.80 - 0.99) | 0.90 (0.79 - 0.98) | 0.85 (0.70 - 0.97) | 0.85 (0.70 - 0.97) |
| Kulkarni_HIV_2 | 0.76 (0.58 - 0.94) | 0.74 (0.52 - 0.91) | 0.73 (0.53 - 0.90) | 0.71 (0.52 - 0.88) |
| Kwan_186 | 0.80 (0.61 - 0.96) | 0.85 (0.60 - 0.97) | 0.80 (0.63 - 0.94) | 0.74 (0.59 - 0.94) |
| LauxdaCosta_OD_3 | 0.86 (0.73 - 0.96) | 0.89 (0.75 - 0.97) | 0.79 (0.61 - 0.96) | 0.83 (0.66 - 0.97) |
| Lee_4 | 0.66 (0.34 - 0.90) | 0.62 (0.37 - 0.89) | 0.63 (0.34 - 0.91) | 0.67 (0.36 - 0.93) |
| Leong_24 | 0.73 (0.60 - 0.92) | 0.76 (0.58 - 0.94) | 0.69 (0.50 - 0.92) | 0.80 (0.61 - 0.97) |
| Leong_RISK_29 | 0.70 (0.39 - 0.97) | 0.74 (0.58 - 0.93) | 0.50 (0.27 - 0.91) | 0.76 (0.56 - 0.95) |
| Long_RES_10 | 0.86 (0.75 - 0.97) | 0.84 (0.71 - 0.96) | 0.84 (0.71 - 0.97) | 0.83 (0.68 - 0.95) |
| Maertzdorf_15 | 0.89 (0.77 - 0.98) | 0.91 (0.79 - 0.99) | 0.83 (0.66 - 0.95) | 0.86 (0.72 - 0.97) |
| Maertzdorf_4 | 0.72 (0.54 - 0.90) | 0.86 (0.75 - 0.98) | 0.71 (0.50 - 0.90) | 0.82 (0.67 - 0.96) |
| Maertzdorf_OD_100 | 0.69 (0.41 - 0.95) | 0.79 (0.55 - 0.97) | 0.73 (0.43 - 0.91) | 0.83 (0.60 - 0.95) |
| Natarajan_7 | 0.85 (0.74 - 0.97) | 0.88 (0.78 - 0.97) | 0.79 (0.63 - 0.96) | 0.85 (0.70 - 0.98) |
| PennNich_RISK_6 | 0.71 (0.56 - 0.92) | 0.91 (0.75 - 0.98) | 0.67 (0.48 - 0.94) | 0.84 (0.70 - 0.97) |
| Qian_OD_17 | 0.87 (0.74 - 0.97) | 0.80 (0.64 - 0.95) | 0.70 (0.55 - 0.95) | 0.72 (0.55 - 0.94) |
| Rajan_HIV_5 | 0.80 (0.64 - 0.95) | 0.75 (0.60 - 0.92) | 0.81 (0.63 - 0.95) | 0.79 (0.63 - 0.94) |
| Roe_3 | 0.85 (0.73 - 0.98) | 0.88 (0.77 - 0.98) | 0.80 (0.68 - 0.96) | 0.82 (0.69 - 0.97) |
| Roe_OD_4 | 0.73 (0.53 - 0.91) | 0.66 (0.51 - 0.89) | 0.70 (0.53 - 0.92) | 0.71 (0.52 - 0.91) |
| Sambarey_HIV_10 | 0.87 (0.71 - 0.96) | 0.83 (0.67 - 0.95) | 0.83 (0.70 - 0.96) | 0.78 (0.67 - 0.95) |
| Singhania_OD_20 | 0.63 (0.39 - 0.94) | 0.86 (0.63 - 0.99) | 0.66 (0.32 - 0.91) | 0.81 (0.60 - 0.97) |
| Sivakumaran_11 | 0.65 (0.38 - 0.88) | 0.83 (0.64 - 0.95) | 0.56 (0.34 - 0.84) | 0.78 (0.63 - 0.96) |
| Suliman_4 | 0.83 (0.69 - 0.95) | 0.90 (0.80 - 0.99) | 0.77 (0.59 - 0.93) | 0.88 (0.74 - 0.98) |
| Suliman_RISK_4 | 0.63 (0.45 - 0.90) | 0.82 (0.65 - 0.98) | 0.66 (0.33 - 0.93) | 0.79 (0.62 - 0.95) |
| Sweeney_OD_3 | 0.91 (0.78 - 0.99) | 0.86 (0.74 - 0.97) | 0.79 (0.65 - 0.96) | 0.83 (0.66 - 0.96) |
| Tabone_OD_11 | 0.84 (0.73 - 0.98) | 0.90 (0.80 - 0.98) | 0.80 (0.67 - 0.95) | 0.82 (0.67 - 0.94) |

Table S4 – *Continued from previous page*
| Signatures | PTB vs. Control |  | PTB vs. LTBI |  |
| --- | --- | --- | --- | --- |
|  | ssGSEA | PLAGE | ssGSEA | PLAGE |
| Tabone.RES_25 | 0.85 (0.76 - 0.98) | 0.85 (0.70 - 0.95) | 0.82 (0.67 - 0.95) | 0.80 (0.65 - 0.94) |
| Tabone.RES_27 | 0.89 (0.78 - 0.97) | 0.86 (0.74 - 0.96) | 0.85 (0.71 - 0.97) | 0.81 (0.68 - 0.96) |
| Thompson_9 | 0.88 (0.76 - 0.97) | 0.81 (0.71 - 0.96) | 0.87 (0.71 - 0.97) | 0.79 (0.67 - 0.95) |
| Thompson.FAIL_13 | 0.76 (0.47 - 0.98) | 0.76 (0.46 - 0.97) | 0.60 (0.38 - 0.91) | 0.63 (0.37 - 0.92) |
| Thompson.RES_5 | 0.64 (0.47 - 0.96) | 0.80 (0.61 - 0.95) | 0.63 (0.44 - 0.92) | 0.71 (0.53 - 0.90) |
| Tornheim_71 | 0.84 (0.72 - 0.98) | 0.73 (0.61 - 0.93) | 0.82 (0.70 - 0.97) | 0.78 (0.68 - 0.98) |
| Tornheim.RES_25 | 0.73 (0.58 - 0.91) | 0.81 (0.57 - 0.94) | 0.83 (0.64 - 0.95) | 0.81 (0.61 - 0.95) |
| Verhagen_10 | 0.53 (0.33 - 0.94) | 0.72 (0.45 - 0.95) | 0.60 (0.33 - 0.91) | 0.60 (0.43 - 0.93) |
| Walter_51 | 0.79 (0.62 - 0.95) | 0.84 (0.67 - 0.96) | 0.74 (0.56 - 0.94) | 0.83 (0.67 - 0.97) |
| Walter.PNA_119 | 0.76 (0.58 - 0.96) | 0.75 (0.58 - 0.95) | 0.71 (0.50 - 0.95) | 0.78 (0.54 - 0.95) |
| Walter.PNA_47 | 0.78 (0.47 - 0.97) | 0.79 (0.58 - 0.92) | 0.73 (0.50 - 0.96) | 0.73 (0.49 - 0.92) |
| Zak.RISK_16 | 0.88 (0.76 - 0.98) | 0.87 (0.74 - 0.98) | 0.84 (0.69 - 0.96) | 0.84 (0.67 - 0.96) |
| Zhao.NANO_6 | 0.49 (0.29 - 0.80) | 0.87 (0.64 - 0.98) | 0.53 (0.31 - 0.91) | 0.77 (0.64 - 0.96) |
| Zimmer.RES_3 | 0.91 (0.79 - 0.99) | 0.86 (0.73 - 0.97) | 0.79 (0.66 - 0.95) | 0.83 (0.66 - 0.95) |

**Table S5:** Weighted mean Specificity and 95% confidence interval for 72 signatures for distinguishing patients with active TB from healthy controls or from LTBI.

| Signatures | PTB vs. Control |  | PTB vs. LTBI |  |
| --- | --- | --- | --- | --- |
|  | ssGSEA | PLAGE | ssGSEA | PLAGE |
| Anderson_42 | 0.72 (0.50 - 0.98) | 0.83 (0.69 - 0.98) | 0.77 (0.49 - 0.95) | 0.87 (0.73 - 0.99) |
| Anderson_OD_51 | 0.84 (0.69 - 0.97) | 0.91 (0.78 - 0.99) | 0.87 (0.71 - 0.98) | 0.89 (0.78 - 1.00) |
| Berry_393 | 0.91 (0.80 - 1.00) | 0.88 (0.77 - 0.98) | 0.88 (0.72 - 0.98) | 0.91 (0.75 - 0.99) |
| Berry_OD_86 | 0.87 (0.72 - 0.98) | 0.89 (0.66 - 0.96) | 0.81 (0.62 - 0.98) | 0.83 (0.68 - 0.95) |
| Blankley_380 | 0.87 (0.78 - 1.00) | 0.90 (0.77 - 0.98) | 0.91 (0.75 - 0.99) | 0.90 (0.78 - 1.00) |
| Blankley_5 | **0.94 (0.83 - 1.00)** | 0.93 (0.81 - 1.00) | 0.89 (0.79 - 1.00) | 0.90 (0.77 - 1.00) |
| Bloom_OD_144 | 0.93 (0.78 - 0.99) | 0.84 (0.73 - 0.98) | **0.92 (0.78 - 0.99)** | 0.86 (0.72 - 0.99) |
| Bloom_RES_268 | 0.93 (0.80 - 1.00) | 0.89 (0.75 - 0.97) | 0.90 (0.77 - 1.00) | 0.88 (0.75 - 0.99) |
| Bloom_RES_558 | 0.88 (0.78 - 0.99) | 0.88 (0.78 - 0.98) | 0.88 (0.74 - 0.99) | 0.90 (0.77 - 0.99) |
| Chen_HIV_4 | 0.87 (0.76 - 0.98) | 0.82 (0.68 - 0.92) | 0.81 (0.68 - 0.97) | 0.84 (0.66 - 0.97) |
| Darboe_RISK_11 | 0.91 (0.78 - 0.99) | 0.89 (0.77 - 0.99) | 0.89 (0.75 - 0.99) | 0.83 (0.72 - 0.99) |
| Dawany_HIV_251 | 0.62 (0.38 - 0.93) | 0.73 (0.54 - 0.92) | 0.66 (0.42 - 0.96) | 0.74 (0.48 - 0.92) |
| Duffy_23 | 0.81 (0.58 - 0.97) | 0.81 (0.64 - 0.93) | 0.81 (0.58 - 0.95) | 0.83 (0.68 - 0.95) |
| Esmail_203 | 0.86 (0.70 - 0.99) | 0.85 (0.73 - 0.98) | 0.79 (0.63 - 0.97) | 0.83 (0.70 - 0.99) |
| Esmail_82 | 0.81 (0.65 - 0.98) | 0.86 (0.70 - 0.95) | 0.82 (0.63 - 0.96) | 0.87 (0.70 - 1.00) |
| Esmail_OD_893 | 0.84 (0.66 - 0.98) | 0.81 (0.67 - 0.95) | 0.85 (0.66 - 0.98) | 0.87 (0.75 - 1.00) |
| Estevez_133 | 0.91 (0.79 - 0.99) | 0.90 (0.77 - 0.99) | 0.91 (0.77 - 0.99) | 0.91 (0.74 - 1.00) |
| Estevez_259 | 0.92 (0.79 - 0.99) | 0.89 (0.68 - 0.96) | 0.90 (0.77 - 0.99) | 0.86 (0.70 - 1.00) |
| Gjoen_10 | 0.80 (0.60 - 0.97) | 0.85 (0.64 - 0.99) | 0.76 (0.48 - 0.93) | 0.84 (0.66 - 0.98) |
| Gjoen_7 | 0.87 (0.66 - 0.97) | 0.86 (0.74 - 0.97) | 0.81 (0.67 - 0.99) | 0.84 (0.72 - 0.98) |
| Gliddon_2_OD_4 | 0.80 (0.61 - 0.98) | 0.88 (0.66 - 0.99) | 0.80 (0.62 - 0.96) | 0.80 (0.62 - 0.96) |
| Gliddon_HIV_3 | 0.92 (0.81 - 0.99) | 0.91 (0.80 - 1.00) | 0.87 (0.71 - 0.98) | 0.88 (0.72 - 1.00) |
| Gliddon_OD_3 | 0.92 (0.81 - 0.99) | 0.91 (0.79 - 1.00) | 0.87 (0.71 - 0.98) | 0.88 (0.72 - 1.00) |
| Gliddon_OD_4 | 0.80 (0.61 - 0.98) | 0.88 (0.66 - 0.99) | 0.80 (0.62 - 0.96) | 0.80 (0.62 - 0.96) |
| Gong_OD_4 | 0.91 (0.81 - 0.99) | 0.89 (0.78 - 0.99) | 0.87 (0.74 - 0.99) | 0.88 (0.74 - 0.99) |
| Heycken_FAIL_22 | 0.92 (0.80 - 0.99) | 0.93 (0.80 - 0.99) | 0.91 (0.76 - 0.99) | 0.89 (0.73 - 1.00) |
| Hoang_OD_13 | 0.92 (0.80 - 1.00) | 0.89 (0.77 - 0.99) | 0.87 (0.73 - 0.99) | 0.83 (0.70 - 0.98) |
| Hoang_OD_20 | **0.94 (0.82 - 1.00)** | 0.90 (0.79 - 0.99) | 0.87 (0.75 - 1.00) | 0.89 (0.73 - 0.99) |
| Huang_OD_13 | 0.84 (0.66 - 0.98) | 0.84 (0.69 - 0.98) | 0.74 (0.58 - 0.96) | 0.88 (0.60 - 0.98) |
| Jacobsen_3 | 0.76 (0.63 - 0.95) | 0.84 (0.72 - 0.98) | 0.77 (0.57 - 0.94) | 0.86 (0.71 - 0.98) |
| Jenum_8 | 0.64 (0.38 - 0.97) | 0.77 (0.61 - 0.94) | 0.72 (0.55 - 0.95) | 0.80 (0.63 - 0.98) |
| Kaforou_27 | 0.92 (0.80 - 0.98) | 0.93 (0.84 - 0.99) | 0.90 (0.76 - 0.99) | 0.90 (0.78 - 1.00) |
| Kaforou_OD_44 | 0.85 (0.66 - 0.96) | 0.87 (0.73 - 0.95) | 0.82 (0.65 - 0.96) | 0.85 (0.72 - 0.99) |
| Kaforou_OD_53 | 0.86 (0.64 - 0.97) | 0.92 (0.84 - 0.99) | 0.80 (0.59 - 0.97) | 0.90 (0.79 - 0.99) |
| Kaul_3 | 0.91 (0.82 - 1.00) | 0.92 (0.81 - 0.99) | 0.89 (0.77 - 0.99) | 0.88 (0.76 - 0.99) |
| Kulkarni_HIV_2 | 0.88 (0.69 - 1.00) | 0.82 (0.65 - 0.98) | 0.84 (0.68 - 0.98) | 0.83 (0.67 - 0.97) |
| Kwan_186 | 0.86 (0.70 - 0.99) | 0.73 (0.61 - 0.95) | 0.87 (0.74 - 0.99) | 0.88 (0.68 - 0.99) |
| LauxdaCosta_OD_3 | 0.90 (0.77 - 0.99) | 0.88 (0.78 - 0.99) | 0.80 (0.63 - 0.97) | 0.86 (0.71 - 0.99) |
| Lee_4 | 0.62 (0.40 - 0.94) | 0.70 (0.45 - 0.93) | 0.71 (0.42 - 0.96) | 0.66 (0.39 - 0.95) |
| Leong_24 | 0.89 (0.71 - 0.99) | 0.79 (0.62 - 0.95) | 0.79 (0.56 - 0.97) | 0.82 (0.65 - 0.97) |
| Leong_RISK_29 | 0.59 (0.35 - 0.92) | 0.82 (0.64 - 0.94) | 0.74 (0.34 - 0.97) | 0.80 (0.61 - 0.97) |
| Long_RES_10 | 0.92 (0.80 - 0.99) | 0.90 (0.78 - 0.99) | 0.91 (0.78 - 0.99) | 0.89 (0.76 - 0.99) |
| Maertzdorf_15 | 0.90 (0.79 - 0.99) | 0.92 (0.82 - 0.99) | 0.85 (0.74 - 0.99) | 0.89 (0.76 - 0.99) |
| Maertzdorf_4 | 0.82 (0.64 - 0.97) | 0.93 (0.80 - 1.00) | 0.77 (0.58 - 0.97) | 0.87 (0.74 - 0.99) |
| Maertzdorf_OD_100 | 0.65 (0.40 - 0.92) | 0.69 (0.49 - 0.91) | 0.69 (0.51 - 0.96) | 0.76 (0.63 - 0.96) |
| Natarajan_7 | 0.90 (0.78 - 0.99) | **0.94 (0.83 - 0.99)** | 0.83 (0.65 - 0.97) | 0.86 (0.73 - 0.99) |
| PennNich_RISK_6 | 0.77 (0.55 - 0.91) | 0.81 (0.72 - 0.94) | 0.74 (0.46 - 0.93) | 0.89 (0.75 - 0.99) |
| Qian_OD_17 | 0.88 (0.76 - 0.99) | 0.86 (0.71 - 0.99) | 0.84 (0.59 - 0.98) | 0.82 (0.58 - 0.97) |
| Rajan_HIV_5 | 0.88 (0.73 - 0.99) | 0.90 (0.76 - 0.99) | 0.86 (0.71 - 0.99) | 0.87 (0.75 - 0.99) |
| Roe_3 | **0.94 (0.81 - 1.00)** | 0.92 (0.81 - 1.00) | 0.91 (0.75 - 0.99) | 0.89 (0.74 - 0.99) |
| Roe_OD_4 | 0.83 (0.67 - 0.98) | 0.88 (0.66 - 0.98) | 0.84 (0.62 - 0.98) | 0.84 (0.66 - 0.99) |
| Sambarey_HIV_10 | 0.86 (0.76 - 0.99) | 0.85 (0.73 - 0.97) | 0.90 (0.77 - 0.99) | 0.93 (0.77 - 0.99) |
| Singhania_OD_20 | 0.69 (0.38 - 0.92) | 0.74 (0.60 - 0.95) | 0.59 (0.33 - 0.94) | 0.77 (0.61 - 0.95) |
| Sivakumaran_11 | 0.70 (0.49 - 0.90) | 0.79 (0.67 - 0.94) | 0.76 (0.50 - 0.96) | 0.88 (0.70 - 0.99) |
| Suliman_4 | 0.89 (0.77 - 0.98) | 0.92 (0.81 - 0.99) | 0.83 (0.66 - 0.97) | 0.88 (0.78 - 0.99) |
| Suliman_RISK_4 | 0.77 (0.52 - 0.94) | 0.82 (0.65 - 0.96) | 0.68 (0.42 - 0.97) | 0.83 (0.65 - 0.95) |
| Sweeney_OD_3 | 0.90 (0.80 - 0.98) | 0.92 (0.82 - 0.99) | 0.91 (0.76 - 1.00) | 0.87 (0.76 - 1.00) |
| Tabone_OD_11 | 0.91 (0.76 - 0.99) | 0.92 (0.83 - 0.99) | 0.91 (0.76 - 0.99) | 0.89 (0.77 - 0.99) |

Table S5 – *Continued from previous page*
| Signatures | PTB vs. Control |  | PTB vs. LTBI |  |
| --- | --- | --- | --- | --- |
|  | ssGSEA | PLAGE | ssGSEA | PLAGE |
| Tabone.RES_25 | 0.93 (0.81 - 0.99) | 0.90 (0.80 - 0.99) | 0.90 (0.75 - 0.98) | 0.89 (0.74 - 0.99) |
| Tabone.RES_27 | **0.94 (0.84 - 1.00)** | 0.92 (0.82 - 1.00) | 0.90 (0.77 - 0.99) | 0.91 (0.77 - 0.99) |
| Thompson_9 | 0.92 (0.83 - 1.00) | **0.94 (0.80 - 0.99)** | 0.89 (0.80 - 1.00) | **0.94 (0.81 - 1.00)** |
| Thompson.FAIL_13 | 0.69 (0.46 - 0.95) | 0.66 (0.43 - 0.94) | 0.77 (0.47 - 0.95) | 0.73 (0.43 - 0.96) |
| Thompson.RES_5 | 0.85 (0.53 - 0.99) | 0.78 (0.63 - 0.95) | 0.81 (0.54 - 0.98) | 0.82 (0.62 - 0.96) |
| Tornheim_71 | 0.92 (0.78 - 0.99) | 0.93 (0.69 - 0.99) | **0.92 (0.77 - 0.99)** | 0.93 (0.74 - 0.99) |
| Tornheim.RES_25 | 0.88 (0.70 - 0.99) | 0.75 (0.62 - 0.96) | 0.83 (0.70 - 0.98) | 0.81 (0.66 - 0.98) |
| Verhagen_10 | 0.75 (0.35 - 0.94) | 0.65 (0.42 - 0.91) | 0.70 (0.39 - 0.95) | 0.79 (0.47 - 0.94) |
| Walter_51 | 0.85 (0.67 - 0.98) | 0.83 (0.70 - 0.96) | 0.84 (0.63 - 0.98) | 0.84 (0.70 - 0.95) |
| Walter.PNA_119 | 0.79 (0.58 - 0.96) | 0.80 (0.60 - 0.96) | 0.76 (0.50 - 0.94) | 0.76 (0.60 - 0.98) |
| Walter.PNA_47 | 0.64 (0.45 - 0.93) | 0.74 (0.60 - 0.94) | 0.68 (0.44 - 0.90) | 0.74 (0.56 - 0.96) |
| Zak.RISK_16 | 0.92 (0.81 - 1.00) | 0.89 (0.76 - 0.99) | 0.89 (0.76 - 1.00) | 0.85 (0.72 - 0.99) |
| Zhao.NANO_6 | 0.83 (0.56 - 0.98) | 0.78 (0.65 - 0.99) | 0.77 (0.39 - 0.96) | 0.89 (0.68 - 0.99) |
| Zimmer.RES_3 | 0.90 (0.80 - 0.98) | 0.92 (0.82 - 0.99) | 0.91 (0.77 - 0.99) | 0.87 (0.76 - 1.00) |

**Figure S1:**
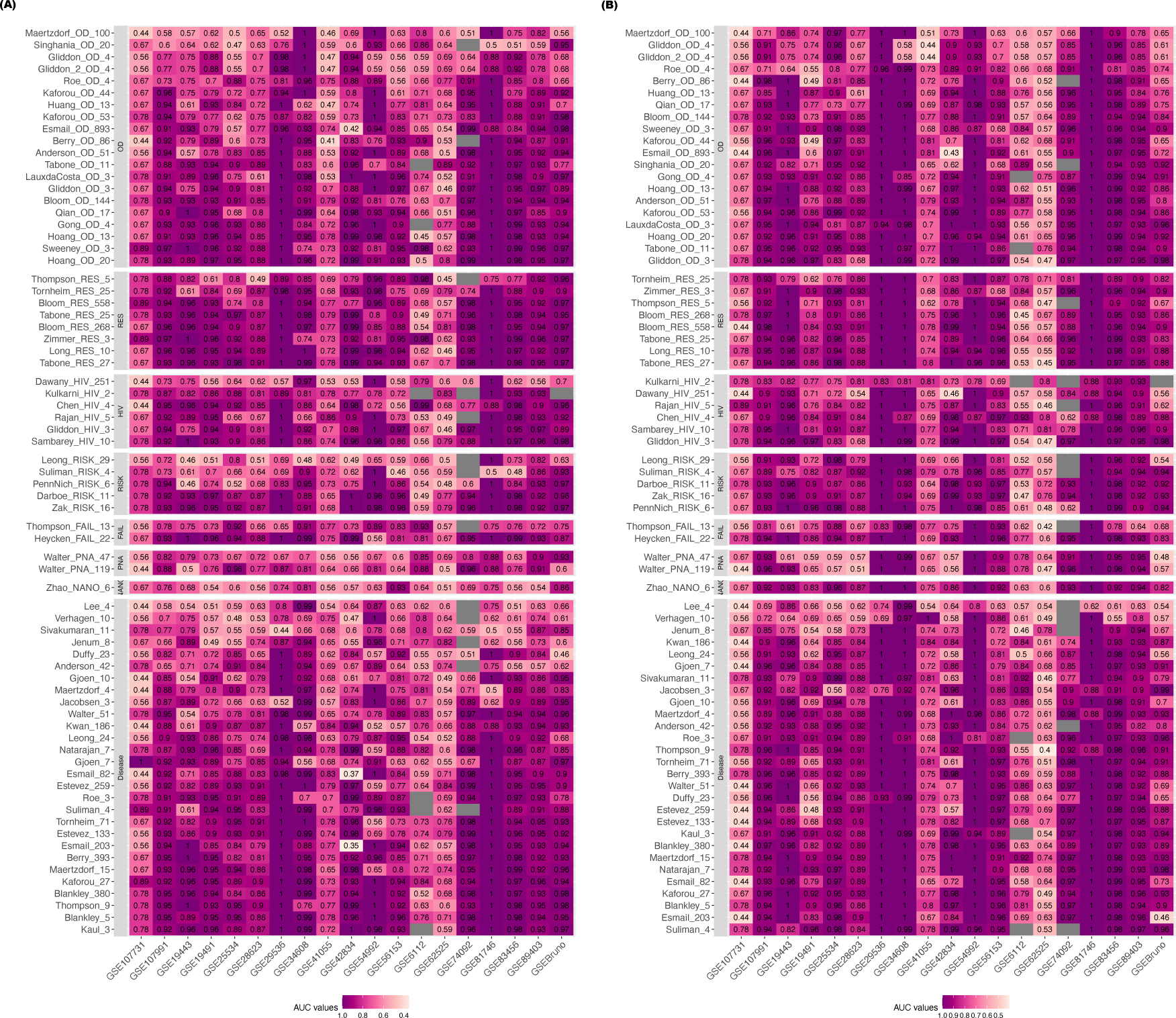
Distribution of AUC values for active TB (PTB) versus healthy controls (control) using ssGSEA **(A)** and PLAGE **(B)**. Gene signatures are clustered based on their subtypes. Gary box indicates that prediction scores was unavailable for the corresponding study.

**Figure S2:**
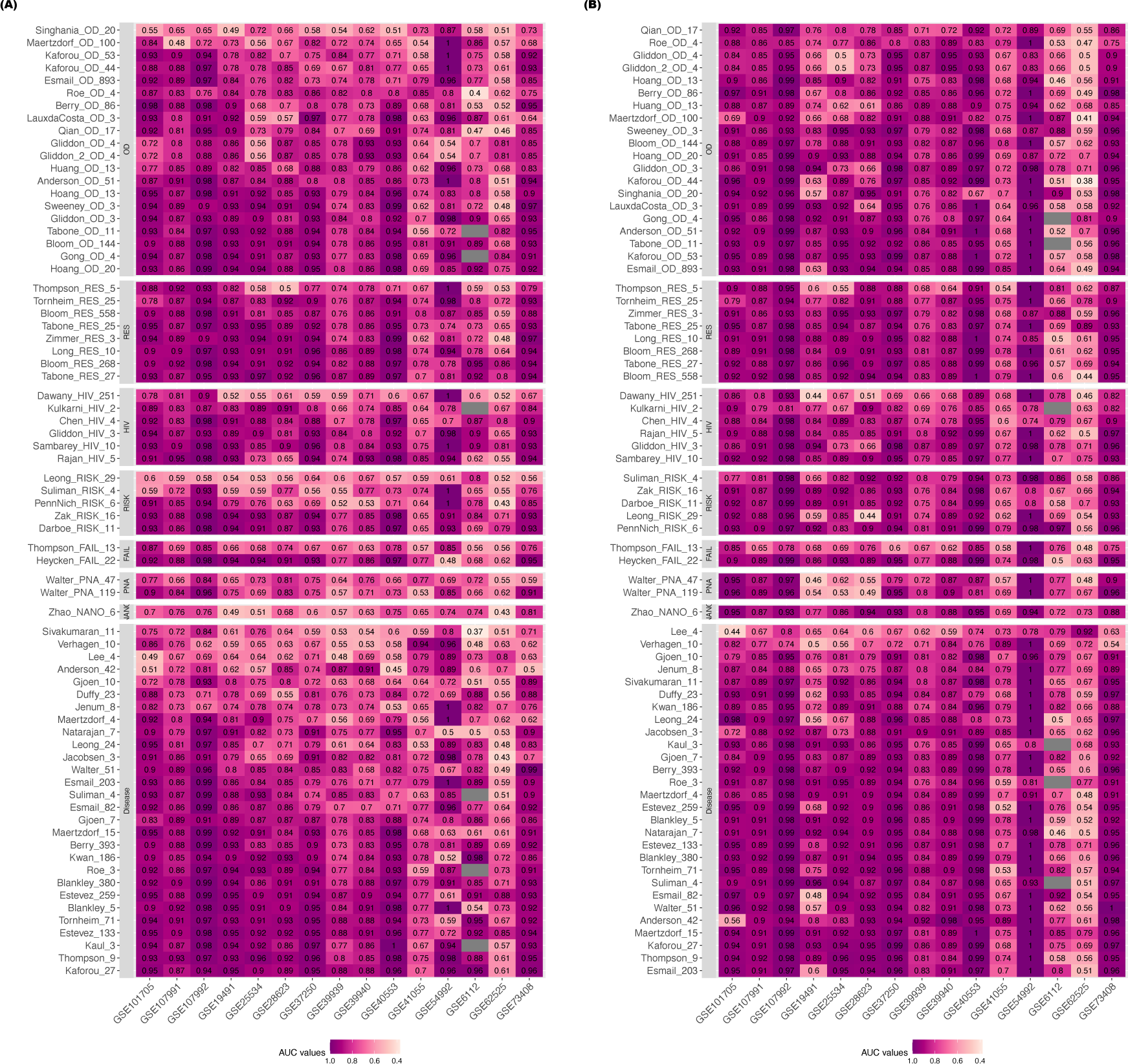
Distribution of AUC values for active TB versus LTBI using ssGSEA **(A)** and PLAGE **(B)**. Gene signatures are clustered based on their subtypes. AUC values were included within each cell, and datasets used to train the signatures are bordered in black. Gary box indicates that signature was not found in the data.

**Figure S3:**
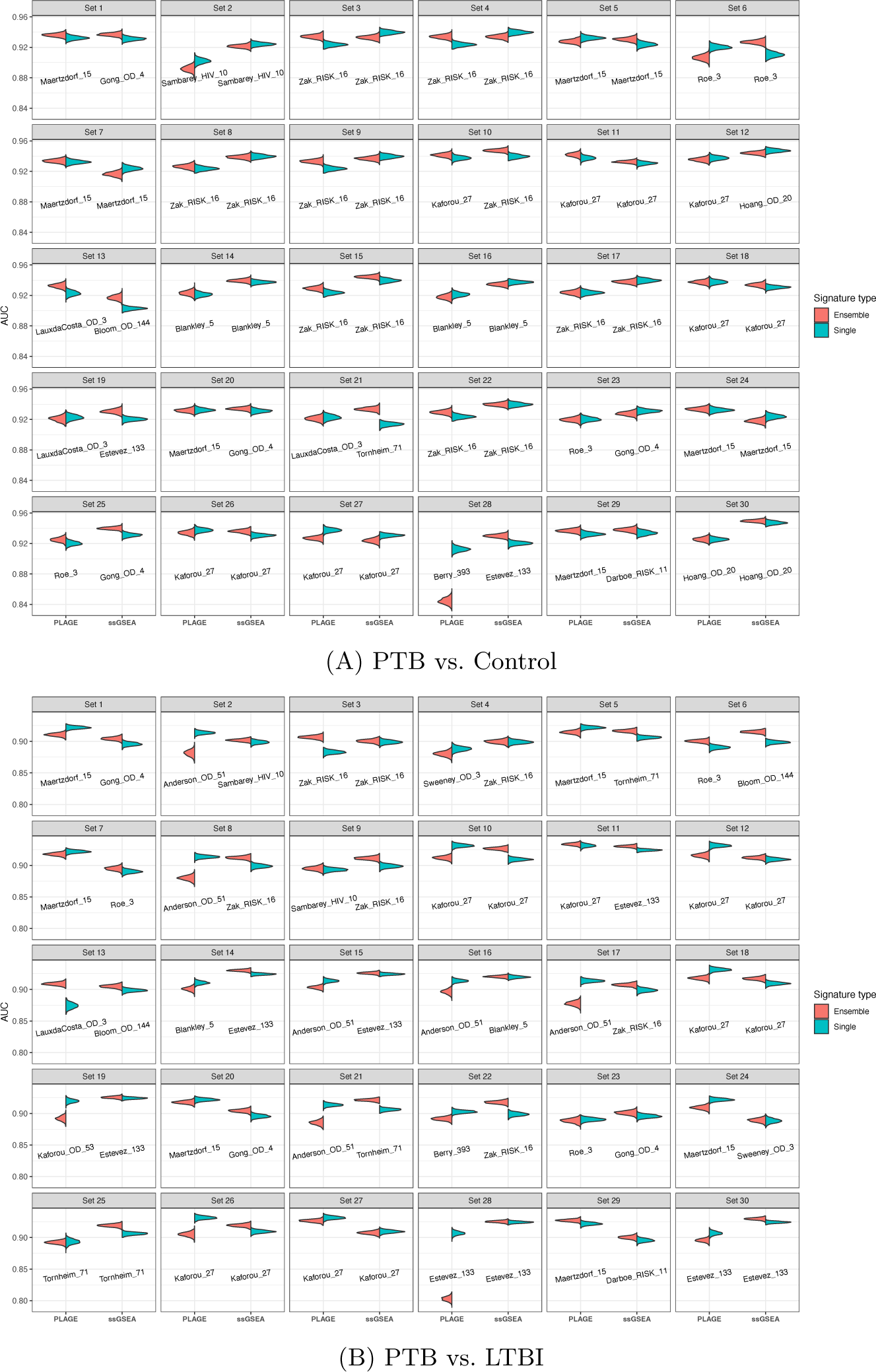
Comparison of 34 ensemble of multiple signatures to each of its single best signature in distinguishing PTB from Control **(A)** or PTB from LTBI **(B)** using PLAGE or ssGSEA. Each set contains 5 TB gene signatures that were randomly selected from 34 TB gene signatures. AUC results were computed based on 300 cross-study validation. Gene signature with the highest weighted mean AUC within the corresponding set was labeled on the plot.

**Figure S4:**
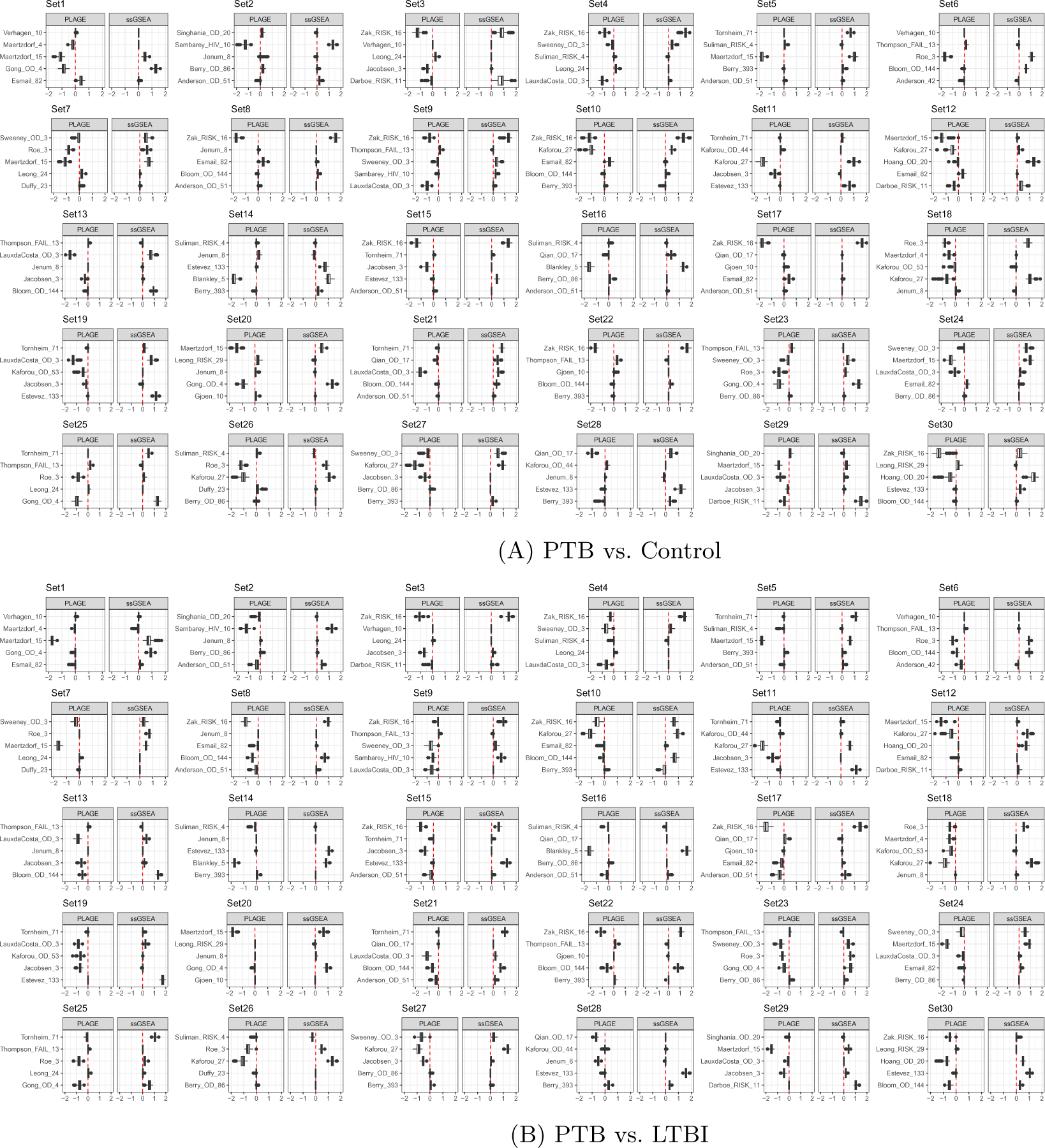
Distribution of coefficients for gene signatures randomly sampled from 34 TB gene signatures for PTB vs. Control **(A)** and PTB vs. LTBI **(B)**. The coefficient distribution was computed based on the results from 300 cross-study validations.

**Figure S5:**
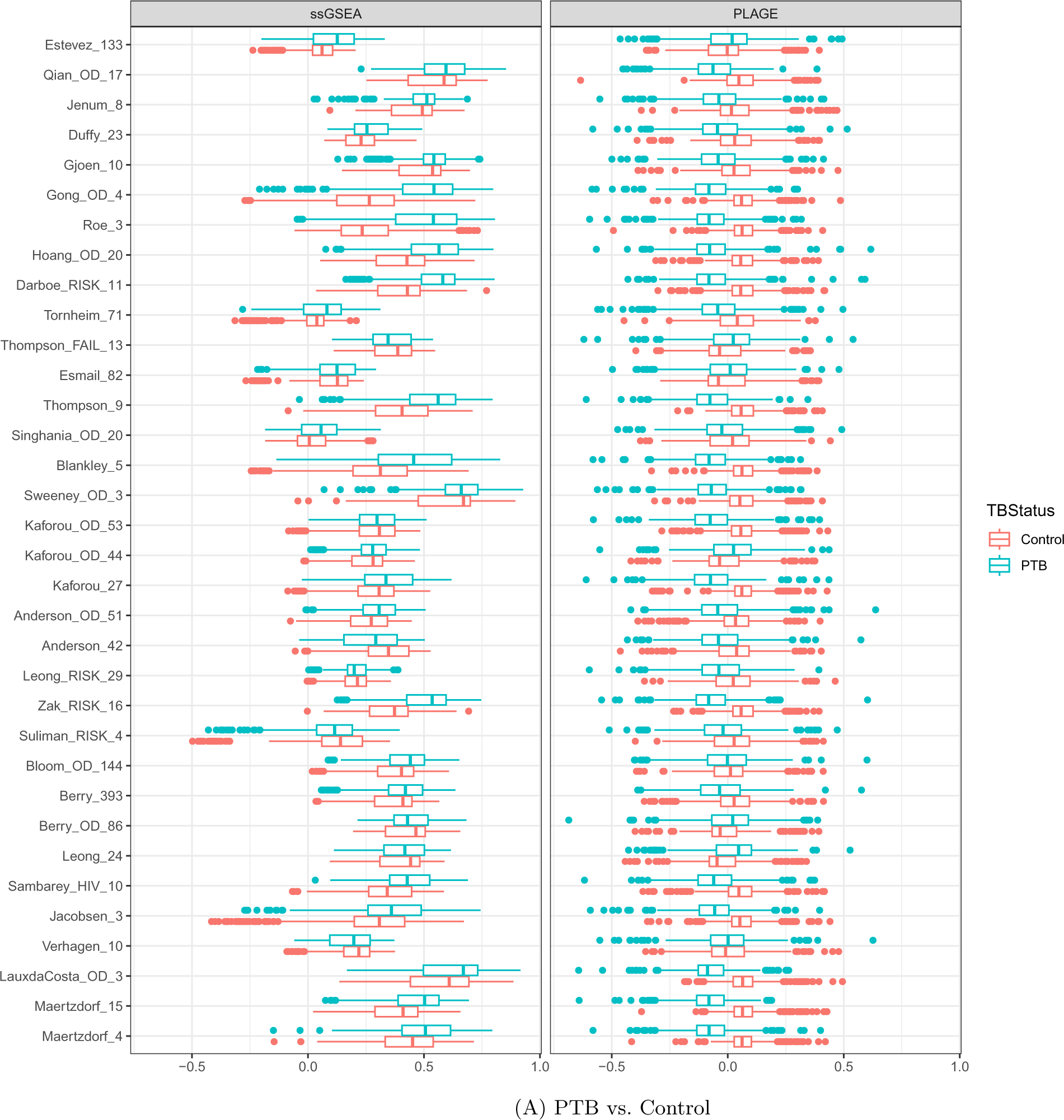
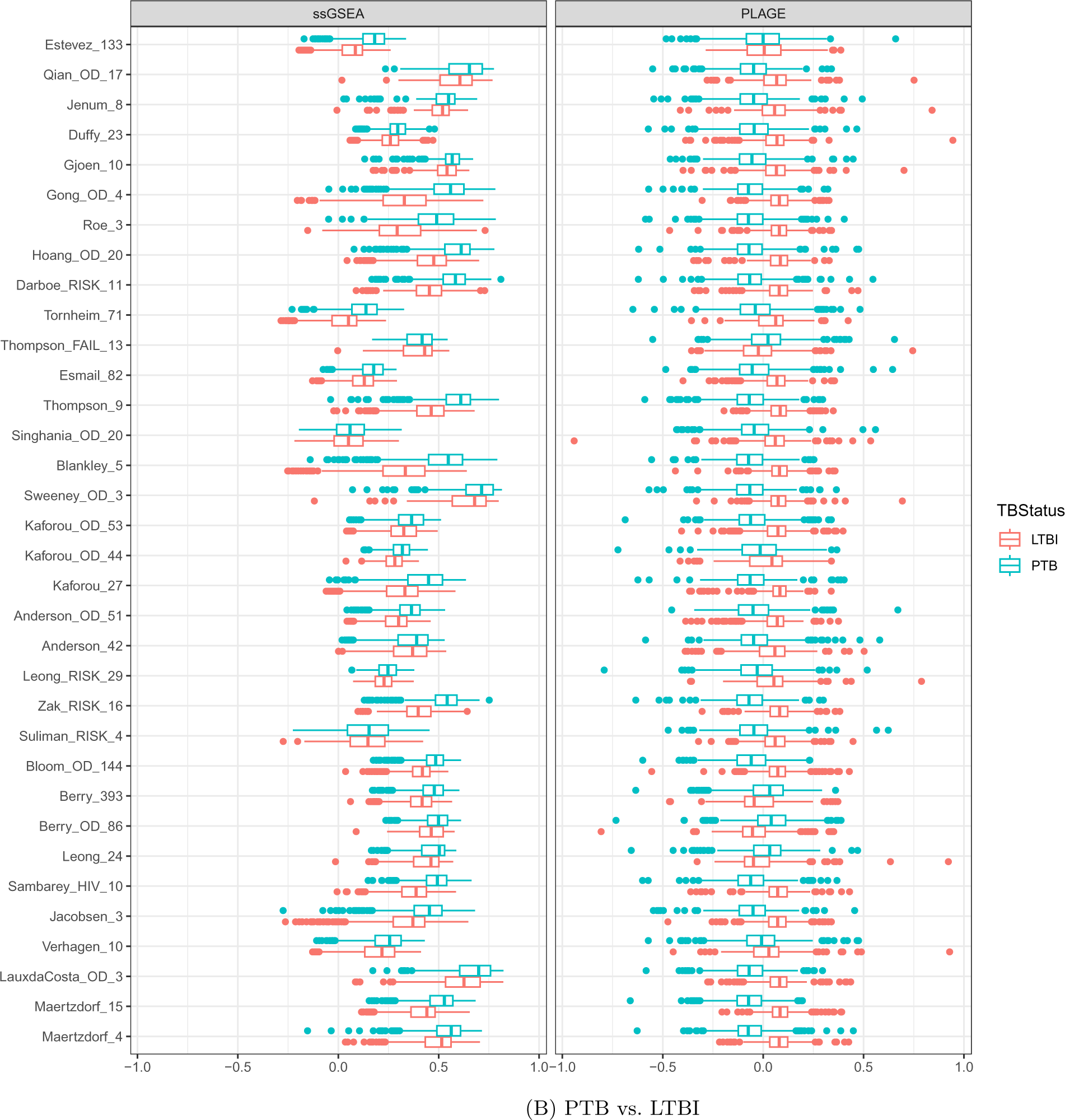
Distribution of profiling scores for 72 signatures given by ssGSEA and PLAGE for PTB vs. Control **(A)** and PTB vs. LTBI **(B)**

**Figure S6:**
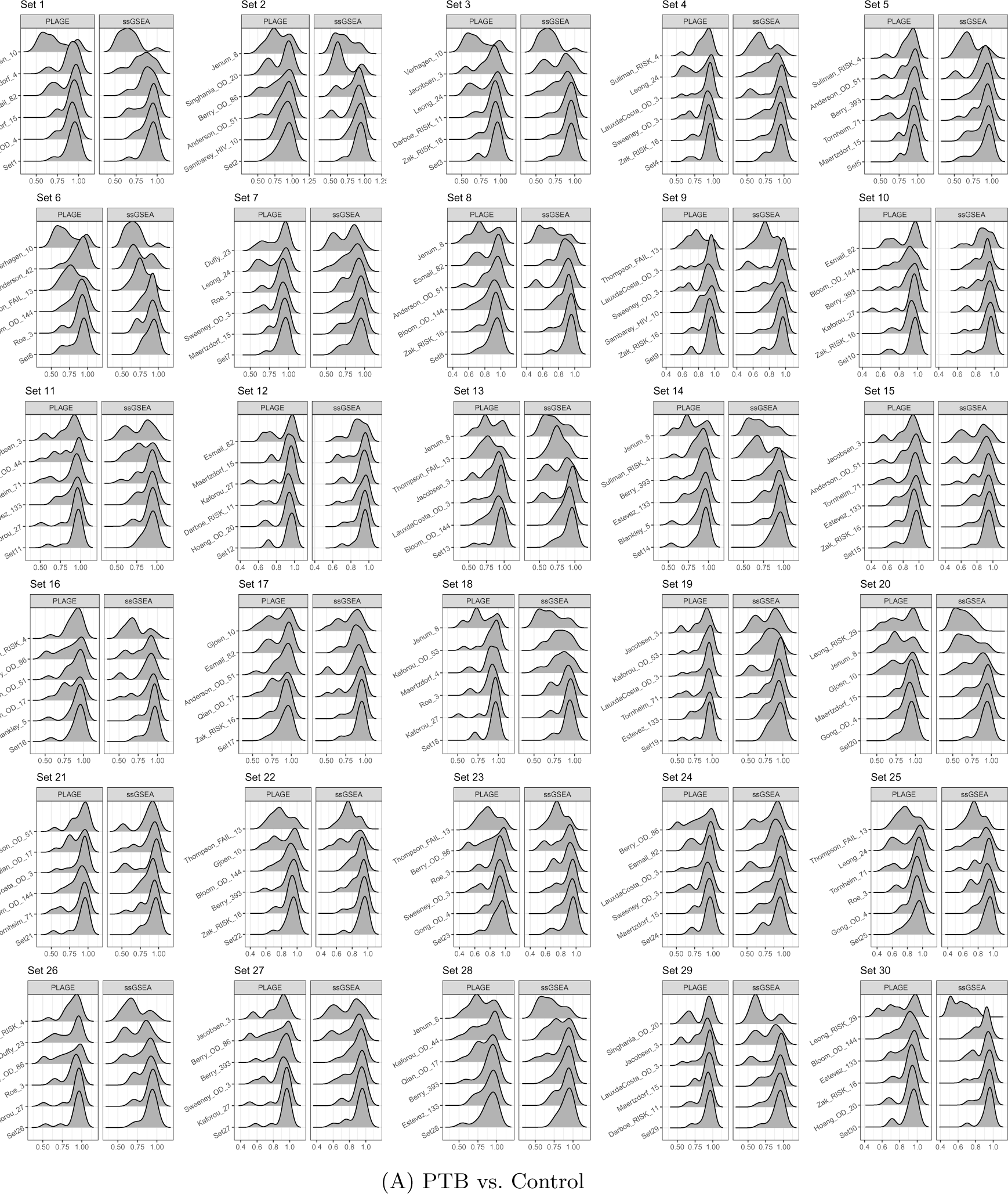
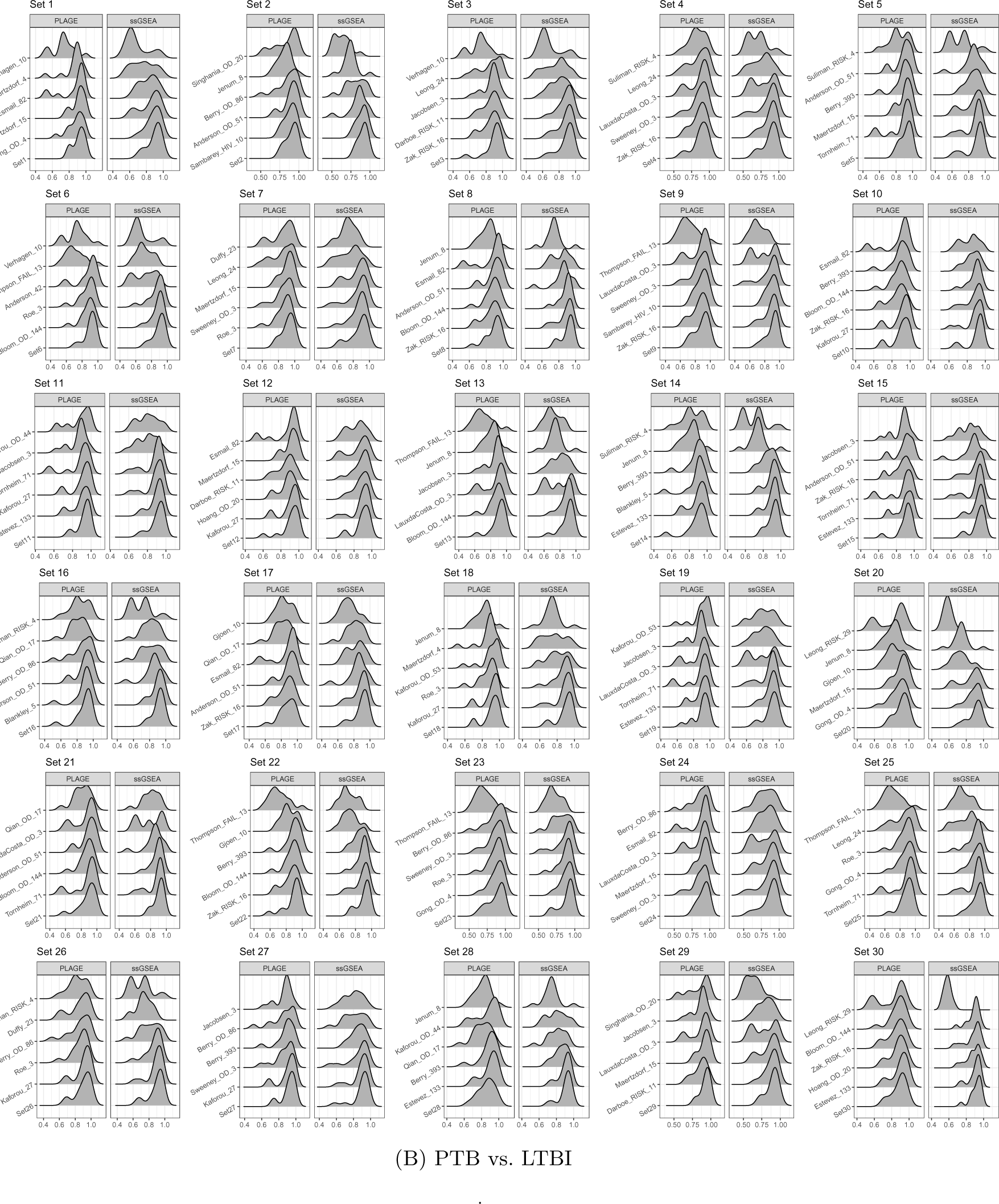
Distribution of AUC values for each ensemble set given by ssGSEA and PLAGE for PTB vs. Control **(A)** and PTB vs. LTBI **(B)**

